# A multiscale model of epigenetic heterogeneity reveals the kinetic routes of pathological cell fate reprogramming

**DOI:** 10.1101/452433

**Authors:** Núria Folguera-Blasco, Rubén Pérez-Carrasco, Elisabet Cuyás, Javier A. Menendez, Tomás Alarcón

## Abstract

The inherent capacity of somatic cells to switch their phenotypic status in response to damage stimuli *in vivo* might have a pivotal role in ageing and cancer. However, how the entry-exit mechanisms of phenotype reprogramming are established remains poorly understood. In an attempt to elucidate such mechanisms, we herein introduce a stochastic model of combined epigenetic regulation (ER)-gene regulatory network (GRN) to study the plastic phenotypic behaviours driven by ER heterogeneity. Furthermore, based on the existence of multiple scales, we formulate a method for stochastic model reduction, from which we derive an efficient hybrid simulation scheme that allows us to deal with such complex systems. Our analysis of the coupled system reveals a regime of tristability in which pluripotent stem-like and differentiated steady-states coexist with a third indecisive state. Crucially, ER heterogeneity of differentiation genes is for the most part responsible for conferring abnormal robustness to pluripotent stem-like states. We then formulate epigenetic heterogeneity-based strategies capable of unlocking and facilitating the transit from differentiation-refractory (pluripotent stem-like) to differentiation-primed epistates. The application of the hybrid numerical method validated the likelihood of such switching involving solely kinetic changes in epigenetic factors. Our results suggest that epigenetic heterogeneity regulates the mechanisms and kinetics of phenotypic robustness of cell fate reprogramming. The occurrence of tunable switches capable of modifying the nature of cell fate reprogramming from pathological to physiological might pave the way for new therapeutic strategies to regulate reparative reprogramming in ageing and cancer.

**Author summary:** Certain modifications of the structure and functioning of the protein/DNA complex called chromatin can allow adult, fully differentiated cells to adopt a stem cell-like pluripotent state in a purely epigenetic manner, not involving changes in the underlying DNA sequence. Such reprogramming-like phenomena may constitute an innate reparative route through which human tissues respond to injury and could also serve as a novel regenerative strategy in human pathological situations in which tissue or organ repair is impaired. However, it should be noted that *in vivo* reprogramming would be capable of maintaining tissue homeostasis provided the acquisition of pluripotency features is strictly transient and accompanied by an accurate replenishment of the specific cell types being lost. Crucially, an excessive reprogramming to pluripotency in the absence of controlled re-differentiation would impair the repair or the replacement of damaged cells, thereby promoting pathological alterations of cell fate. A mechanistic understanding of how the degree of chromatin *plasticity* dictates the reparative versus pathological behaviour of in vivo reprogramming to *rejuvenate* aged tissues while preventing tumorigenesis is urgently needed, including especially the intrinsic epigenetic heterogeneity of the tissue resident cells being reprogrammed. We here introduce a novel method that mathematically captures how epigenetic heterogeneity is actually the driving force that governs the routes and kinetics to entry into and exit from a pathological pluripotent-like state. Moreover, our approach computationally validates the likelihood of unlocking chronic, unrestrained pluripotent states and drive their differentiation down the correct path by solely manipulating the intensity and direction of few epigenetic control switches. Our approach could inspire new therapeutic approaches based on *in vivo* cell reprogramming for efficient tissue regeneration and rejuvenation and cancer treatment.

## Introduction

The correlation between ageing and cancer incidence rate is a well established empirical fact. The currently accepted explanation for such a correlation is subsumed under the multiple hit hypothesis or Knudson hypothesis [1,2]. This multi-mutation theory considers cancer as the result of the successive accumulation of genetic mutations and, therefore, time is necessary for cells to overcome a certain mutagenic threshold before cancer can develop. An alternative paradigm has emerged in recent years, in which the ability of the ageing process per se to interfere with the robustness of the epigenetic regulation (ER) of differentiated phenotypes might suffice to generate malignant transformation [3].

Fully committed somatic cells can spontaneously reprogram to pluripotent stem-like cells during the normal response to injury or damage *in vivo* [4]. Such cellular processes involving dedifferentiation and cell-fate switching might constitute a fundamental element of a tissue’s capacity to self-repair and rejuvenate [5,6]. However, such physiological (normal) cell reprogramming might have pathological consequences if the acquisition of epigenetic and phenotypic plasticity is not transient. In response to chronically permissive tissue environments for *in vivo* reprogramming, the occurrence of unrestrained epigenetic plasticity might permanently lock cells into self-renewing pluripotent cell states disabled for reparative differentiation and prone to malignant transformation (see Fig. 1) [3,7–10].

**Fig 1.**
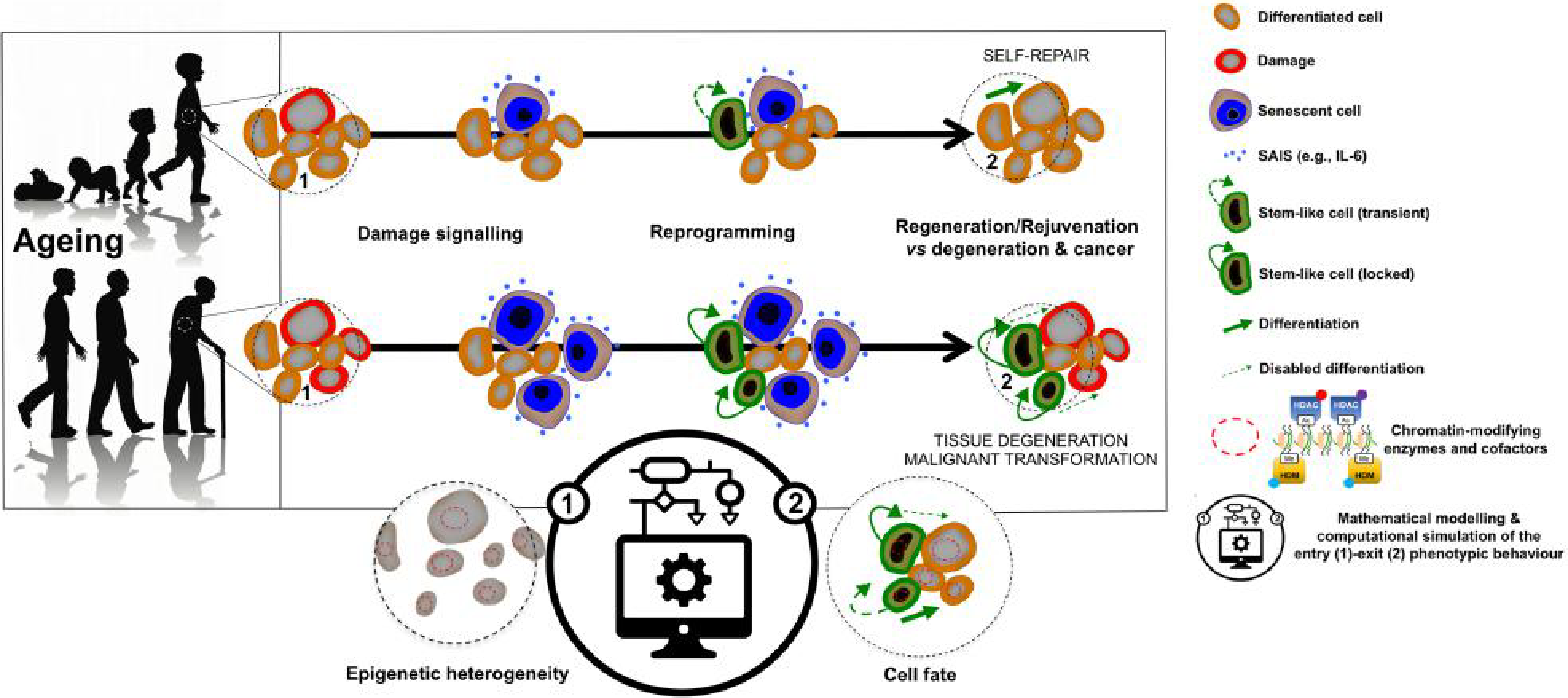
Physiological and pathological cell fate reprogramming: A mathematical approach. Reprogramming-like phenomena in response to damage signalling may constitute a reparative route through which human tissues respond to injury, stress, and disease via induction of a transient acquisition of epigenetic plasticity and phenotype malleability. However, tissue regeneration/rejuvenation should involve not only the transient epigenetic reprogramming of differentiated cells, but also the committed re-acquisition of the original or alternative committed cell fate. Chronic or unrestrained epigenetic plasticity would drive ageing/cancer phenotypes by impairing the repair or the replacement of damaged cells; such uncontrolled phenomena of *in vivo* reprogramming might also generate cancer-like cellular states. Accordingly, we now know that chronic senescence-associated inflammatory signalling might lock cells in highly plastic epigenetic states disabled for reparative differentiation and prone to malignant transformation. We herein introduce a first-in-class stochastic, multiscale reduction method of combined epigenetic regulation (ER)-gene regulatory network (GRN) to mathematically model and computationally simulate how ER heterogeneity regulates the entry-exit mechanisms and kinetics of physiological and pathological cell fate reprogramming. (SAIS: Senescence-associated inflammatory signalling).

Central to such so-called stem-lock model for ageing-driven cancer [8,11,12] is the notion that ER of cell fate should be key for the causal relationship between ageing and cancer. ER refers to a series of modifications of the cell’s DNA without modifying its genetic sequence. Such modifications can disrupt or allow expression of particular genes. By switching on or off different parts of the genome, ER is in fact responsible for the variety of phenotypes in complex multicellular organisms (where all somatic cells are genetically identical). Recent advances in experimental determination of ER mechanisms have triggered an interest in developing mathematical models regarding both ER of gene expression [13–18] and epigenetic memory [14–16,19–22].

We are rapidly amassing evidence that, beyond the role of genetic alterations, non-genetic stimuli such as inflammation or ageing, among others, can promote *epigenetic plasticity*, namely, overly restrictive epigenetic states - capable of preventing the induction of tumour suppression programmes or blocking normal differentiation – or overly plastic epigenetic states – capable of stochastically activating oncogenic programmes and non-physiological cell fate transitions including those leading to the acquisition of stem cell-like states [18,23]. Disruption of normal epigenetic resilience has the potential to give rise to each classic cancer hallmark [23]. An ideal therapy should not only need to address chronic epigenetic plasticity of senescence-damaged tissues, but also to “unlock” stem cell-like states to drive tissue regeneration. Unfortunately, most current models and drug discovery strategies are largely biased on the prevailing, mutation theory of the origin of cancer. Such a framework cannot capture the stochastic aspects that drive the structure and dynamics of epigenetic plasticity and chromatin structure that connects ageing and cancer, and might arise in the absence of initiating genetic events.

Identification of the molecular interactions controlling the transition between normal, restrictive and permissive chromatin states is expected to have major impact in the understanding and therapeutic management of the causative relationship between ageing and cancer [9,24,25]. In this regard, our mathematical approach intends to deconstruct and model the predictive power that epigenetic landscapes might have for the susceptibility of cells to lose their normal identity. In order to determine the key mechanisms underlying epigenetic plasticity and its connections with stem cell-locking, we consider a gene network model that regulates the phenotypic switch between differentiated and pluripotent states. Each gene within this regulatory system is acted upon by epigenetic regulation which restricts/enables its expression capability (see Fig. 2(a) for a schematic representation). Crucially, the precise deconstruction of the highly complex mechanisms through which epigenetic plasticity allows cells to stochastically activate alternative regulatory programs and undergo pathological versus physiological cell fate transitions requires the incorporation of a central role for epigenetic heterogeneity in phenotypic plasticity [18,26] (see Fig. 1 for a schematic representation). Pour et al. [26] have reported evidence that the potential to reprogram is higher within select subpopulations of cells and that preexisting epigenetic heterogeneity can be tuned to make cells more responsive to reprogramming factor induction. Along this line, we recently presented a model of cell fate reprogramming, in which the heterogeneity of epigenetic metabolites, which operates as a regulator of the kinetic parameters promoting/preventing histone modification, stochastically drives phenotypic variability capable of producing cell epistates primed for reprogramming [18]. However, how the variability of both histone-modifying enzymes and their corresponding cofactors might dictate changes in chromatin-related barriers and lead to drastic changes in the transcriptional regulatory networks for cell fate determination was not contemplated.

**Fig 2.**
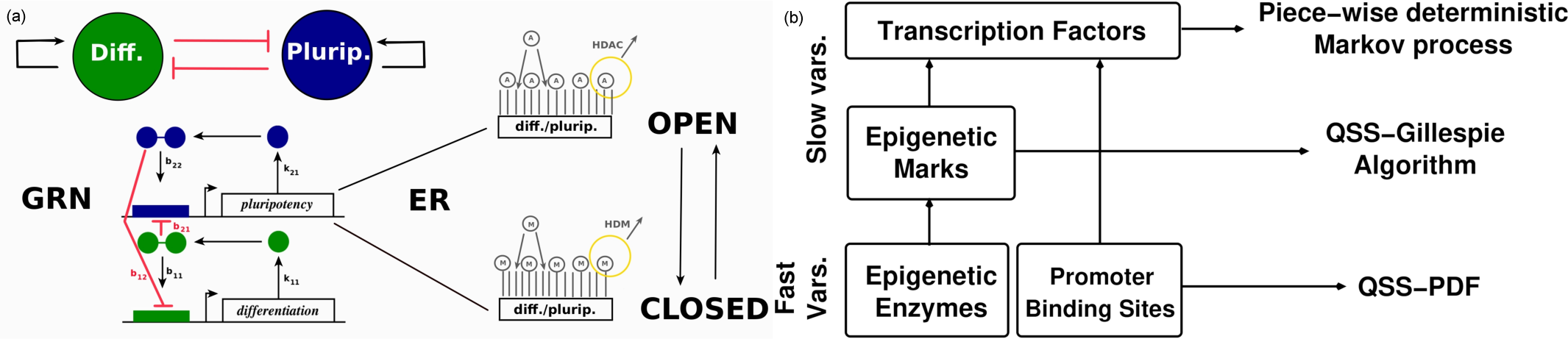
Schematic reprentation of the ER-GRN model and its multiscale reduction. (a): Gene regulatory network (GRN) of two self-activating, mutually-inhibitory genes with epigenetic regulation. In the GRN model, the gene product (denoted by *X_i_* in S2 Table) is its own transcription factor which, upon dimerisation, binds the promoter region of the gene thus triggering gene transcription. The transition rates corresponding to this GRN are given in S2 Table. For simplicity, we use an effective model in which the formation of the dimer and binding to the promoter region is taken into account in a single reaction, and the resulting number of promoter sites bound by two transcription factors is denoted by *X_ij_* (see S2 Table). Furthermore, depending on whether the epigenetic state is open (i.e. predominantly acetylated (A)) or closed (i.e. predominantly methylated (M)) the promoter region of the gene is accessible or inaccessible to the transcription factor, respectively. (b): Schematic representation of the time separation structure of the multiscale method developed to simulate the ER-GRN system. See text and S1 File for more details.

In the present work, we present a stochastic model of a coupled ER-GRN system aimed at analysing the effects of epigenetic plasticity on cell-fate determination and reprogramming driven by the heterogeneity of the ER system. Furthermore, we introduce a stochastic model reduction analysis based on multiple scale asymptotics of the combined ER-GRN system [27–34], which allows us to study a variety of different behaviours due to heterogeneity in the epigenetic regulatory system [26]. By adding ER to the picture, this work extends previous approaches where phenotypes are associated with the attractors of complex gene regulatory systems and their robustness, with the resilience of such attractors in the presence of intrinsic noise, environmental fluctuations, and other disturbances [35–43]. We initially evaluate the epigenetic parameters regulating the entry into robust epigenetic states throughout the entire ER-GRN system. We then formulate epigenetic heterogeneity-based strategies capable of directing the exit and transit from stem-locked to differentiation-primed epistates. We finally apply a hybrid numerical method derived from our theoretical analysis to determine the efficiency of the epigenetic strategies formulated to unlock a persistent state of pathological pluripotency.

This work is organised as follows. In section *Materials and methods*, we present a summary of the formulation of our ER-GRN model and its analysis. By developing a multiscale asymptotic theory to study ER-GRN systems (for which additional details are given in the Supplemental Information), we are able to reduce the complexity of the model to a hybrid system, for which a numerical simulation method was implemented. *Results* section is devoted to a detailed presentation of our results, especially those pertaining to the mechanisms regulating the phenotypic robustness of pluripotency-locked and differentiation-primed states. In the *Conclusion* section, we summarise our findings and present our conclusions.

## Materials and methods: Model formulation and analysis

In this paper, we aim to study an ER-GRN model which can describe cell differentiation and cell reprogramming. One of the simplest GRNs which allows to do this consists of two genes, one promoting differentiation, and the other promoting pluripotency (see Fig. 2(a)). Nevertheless, in this section, we formulate our model and we analyse it considering the most generic case, i.e. we assume to have an arbitrary number of genes *N_G_*. By doing so, our theoretical analysis can be further applied to any ER-GRN model, which implies a wide applicability of the derived formulation. However, when possible, we try to relate the theory developed to our particular ER-GRN so as to keep track of our case study.

### General description of the stochastic model of an epigenetically-regulated gene network

Consider a gene regulatory network composed of *N_G_* self-activating genes which can repress each other. In particular, we consider that the gene product of each of these genes forms homodimers, which act as a transcription factor (TF) for its own gene by binding to its own promoter. Furthermore, each gene within the network has a number of inhibitors, which operate via competitive inhibition: the homodimers of protein *j* bind to the promoter of gene *i*, and by doing so they impede access of the TF to the promoter of gene *i*. In Fig. 2(a), an illustrative scheme of the simplified case of two mutually inhibiting genes, one promoting pluripotency (blue) and one promoting differentiation (green), is shown. The regulation topology of the network can be represented using a weighted adjacency matrix B. B is a *N_G_* × *N_G_* matrix, whose elements, *b_ij_* > 0, are the binding rates of homodimers of protein *j* to the promoter of gene *i* (see Fig. 2(a)). Moreover, the expression of gene *i* is induced at a constant basal production rate, *ℛ_i_*, independent of the regulatory mechanism described above. Proteins (TF monomers) of type i are synthesised at a rate proportional to the number of bound promoter sites with rate constant *k_i1_* and degraded with degradation rate *k_i2_* (see Fig. 2(a), S1 Table and S2 Table).

In addition to TF regulation, we further consider that each gene is under epigenetic regulation (ER). ER controls gene transcription by modulating access of TFs to the promoter regions of the genes. In other words, in our model, ER is associated with an upstream drive that regulates gene expression [44]. Such epigenetic control is often related to alternative covalent modifications of histones. To address the high complexity of ER, we focus on a simpler stochastic model of ER, based on that formulated in [13,18] and [19]. Our model belongs to a wider class of models which consider that single unmodified (U) chromatin loci can be modified so as to acquire positive (A) or negative (M) marks. These positive and negative marks involve covalent modification of histones. Of such modifications we consider methylation (associated with negative marks) and acetylation (associated with positive marks) [19]. An illustrative example on how epigenetic modifications, acetylation and methylation, alter the accessibility of TFs to the promoter regions of the genes is shown in Fig. 2(a). Both modifications are mediated by associated enzymes: histone methylases (HMs) and demethylases (HDMs), and histone acetylases (HACs) and deacetylases (HDACs). For simplicity, we only explicitly account for HDM and HDAC activity (see Fig. 2(a)). In our model, a positive feedback mechanism is introduced whereby M marks help to both add more M marks and remove A marks from neighbouring loci. The positive marks are assumed to be under the effects of a similar positive reinforcement mechanism [16,19]. A full description of the details of the ER model are given in Section S.2 of the S1 File (see also [18]) and S3 Table, where the transition rates for the ER model are given.

Under suitable conditions, determined by the activity and abundance of histone-modifying enzymes and co-factors, the positive reinforcement mechanism produces robust bistable behaviour. In this bistable regime, the two possible ER stable states are: an *open* epigenetic state where the levels of positive (negative) marks are elevated (downregulated). In this case, the promoter of the gene is accessible to TFs and transcription can occur. By contrast, in the absence (abundance) of positive (negative) marks the gene is considered to be *silenced*, as TFs cannot reach the promoter.

An essential part of the stochastic dynamics of the ER system is the noise-induced transitions between the open and silenced states. Escape from steady states is a well-established phenomenon (see e.g. [45]) and thoroughly analysed within the theory of rate processes [46] and large deviation theory [40,47,48]. As we will illustrate below, these noise-induced dynamics are essential to classify the epiphenotypes of somatic cells [18] and stem cells and unravel the mechanisms of reprogramming and locking.

### Multi-scale analysis and model reduction

The system that results from coupling the ER and GRN models becomes rather cumbersome and computationally intractable as the GRN grows. For this reason, in order to analyse the behaviour of the resulting stochastic model, we take advantage of intrinsic separation of time scales [27–34]. We exploit this time scale separation to reduce our model by performing stochastic quasi-steady state approximations (QSSA) by means of asymptotic analysis of the stochastic ER-GRN system as established in [29–31,33] (see Fig. 2(b)). Especifically, we assume that the characteristic scale for the number of TF monomers (*S*), the number of promoter binding sites (*E*), the number of ER modification sites (*Y*), and the number of ER enzymes (*Z*), are such that *S* ≫ *E*, *Y* ≫ *Z* and *O*(*E*) = *O*(*Y*) (see S1 Table for the definition of these variables). Note that the assumption *Y* ≫ *Z* is exactly the Briggs-Haldane hypothesis for enzyme kinetics [49] since the ER modification sites are the substrates for the ER enzymes (see Section S.2 in the S1 File). The multiscale analysis is carried out in this Section, with additional technical details provided in Sections S.3 and S.4 of the S1 File. Furthermore, the corresponding numerical method is described in S1 Appendix. We show that, upon appropriate assumptions regarding the characteristic scales of the different molecular IQS species, our model exhibits a hierarchy of time scales, which allows to simplify the model and its computational simulation.

The resulting reduced stochastic model is such that, since *S* » *E* and *Y* » *Z*, the number of bound-to-promoter TFs and ER enzyme-substrate complexes are fast variables that can be sampled from their quasi-equilibrium distribution with respect (or conditioned to) their associated slow variables. TFs and ER modification sites (ER substrates) are slow variables whose dynamics, which dominate the long time behaviour of the system, are given by their associated stochastic dynamics with the fast variables sampled from their quasi-steady state approximation (QSSA) probability density functions (PDFs). The assumption that *S* ≫ *Y* allows for further simplification of the 20s model, as it allows to take the limit of *S* » 1 in the stochastic equations for the TFs monomers which leads to a *piece-wise deterministic* Markov description: the dynamics of the number of TFs monomers is given by an ODE which is perturbed at discrete times by a noise source [31].

We present a summarised version of the asymptotic model reduction. Details of this analysis are provided in Section S.3 of the S1 File. The starting point of our analysis is the so-called Poisson representation of the stochastic process, which is equivalent to the Master Equation, [30]:

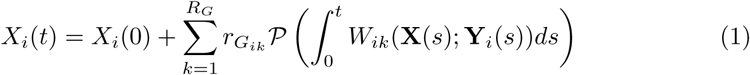

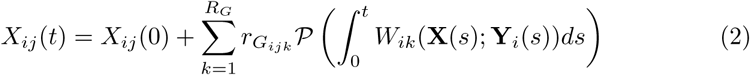

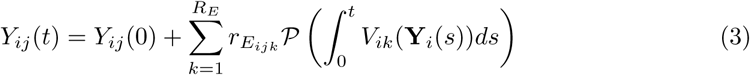

where *X_i_* denotes the product of gene *i*, *X_ij_* refers to the number of dimers of type *j* bound to the promoter region of gene *i* and *Y_j_* corresponds to the number of molecular species of type *j* within the ER model of gene *i*. We also use the notation 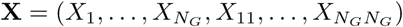 and Y*_i_* = (*Y_i1_*,…,*Y_i7_*). Pλ ∿ Poisson(λ), i.e. P(λ) is a random number sampled from a Poisson distribution with parameter λ [30], *R_G_* and *R_E_* denote the total number of reactions in the GRN model (see S2 Table) and in the ER model (S3 Table), respectively, with *W_ik_* and *V_ik_* denoting the transition rates corresponding to the GRN model and the ER model (see S2 Table and S3 Table, respectively). The stoichiometries 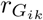, 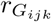 and 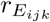 denote the change in number of molecules that reaction *k* has on *X_i_*, *X_ij_* and *Y_ij_*, respectively. Eqs. (1) and (2) are associated with the the stochastic dynamics of the GRN (see S2 Table), which are regulated by the ER part of the model. Eq. (3) describes the dynamics of the ER system, which drives the dynamics of the GRN (see S2 Table and S3 Table).

Under the appropriate conditions, separation of time scales can be made explicit by re-scaling the random variables and the transition rates. Based on our previous work [18,32,34], we propose the following rescaling:

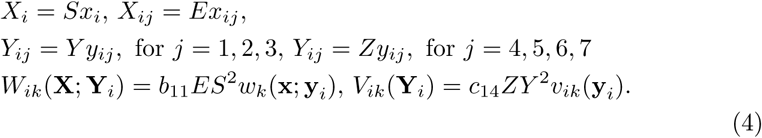

In Eq. (4), the scale factors *S*, *E*, *Y*, and *Z* are the characteristic number of protein transcripts, promoter region binding sites, histone modification sites, and epigenetic enzymes (HDMs and HDACs), respectively. For simplicity, we assume that these scales are the same for all the genes involved in the GRN. We assume that *S* ≫ *E* ≈ *Y* ≫ *Z*. We further define a re-scaled (dimensionless) time: τ = *b_11_ESt*.

By using Eq. (4) in Eqs. (1)-(3), we obtain:

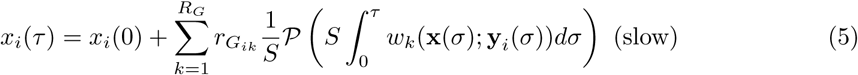

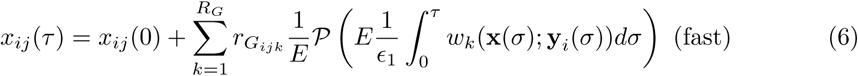

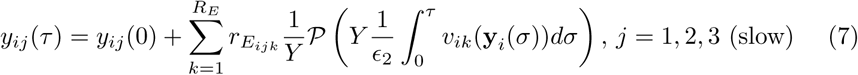

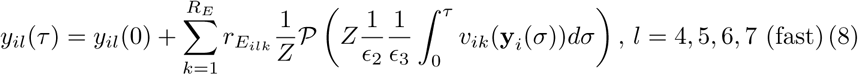

where 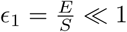, 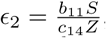, and 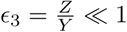, with *∊*_1_ < *∊*_3_. We have no direct information to estimate the order of magnitude of *∊*_2_. Thus, without loss of generality we will assume that *∊*_2_ = *O*(1).

The scaling hypothesis *S* ≫ *E* ≈ *Y* ≫ *Z* allows for a series of successive approximations which enables us to reduce the model Eqs. (5)-(8) into a much less computationally demanding system. First, provided that both ∊_1_ ≫ 1 and ∊_3_ ≫ 1, we can assume that the (rescaled) rates associated with the fast variables GRN-ER dynamics (Eqs. (6) and (8)) are much larger than those corresponding to their slow counterparts (Eqs. (5) and (7)). Under these conditions, the stochastic dynamics of the fast variables reaches their (quasi-)steady states while the slow variables are effectively frozen [27–29,50].

We proceed with the asymptotic model reduction by first addressing the QSSA PDFs of the fast variables (see *Inner solution* below). We then move on to study the QSS approximation of the slow variables, in particular, the large-*S* asymptotics of the protein concentration dynamics (see *Outer solution* below).

#### Inner solution

The inner solution corresponds to the relaxation dynamics of the fast variables onto their quasi-equilibrium state, while the slow variables remain unchanged. The solution of the inner dynamics allows us to determine the QSSA PDFs of the fast variables conditioned to fixed values of the slow variables.

We proceed by considering the following rescaling of the time variable 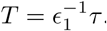. Upon such rescaling, it is straightforward that all the rates of the reactions affecting the slow variables (Eqs. (5) and (7)) are now *O*(∊_1_), which implies that the slow variables, *x_i_* and *y_ij_* (for *j* = 1, 2, 3), can be considered to remain frozen whilst the fast variables reach their quasi-equilibrium distribution according to the dynamics:

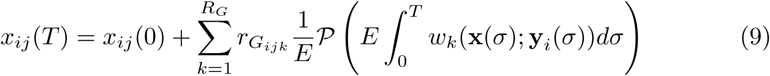

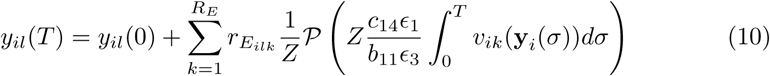

where *l* = 4, 5, 6, 7, *∊*_2_ = *O*(1) and the slow variables, *x_i_* and *y_ij_*, *j* = 1, 2, 3, are considered to stay constant.

Consider the (inner) dynamics of the number of bound sites within the promoter regions, *X_ij_*(*T*), Eq. (9). Provided that the ER of gene *i* remains in the open state, i.e. *y_i2_* ≪ 1 and *y_i3_* ∿ O(1), the resulting stochastic dynamics describes how the binding sites switch between bound-to-TF dimer to unbound-to-TF dimer at constant rates (since the number of the different TF molecules does not change at this time scale). Since the number of binding sites is a constant, the (quasi-)steady state distribution of bound TFs to each promoter is a multinomial (see Section 3.2 of the S1 File for a detailed derivation of this result). Otherwise, if gene *i* is epigenetically closed, then *X_ij_* (*T*) = 0 for all *j* with probability one. Therefore, the random vector describing the number of TFs bound at the promoter region of gene *i*, **B***_i_*, whose components are 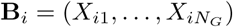 is sampled from:

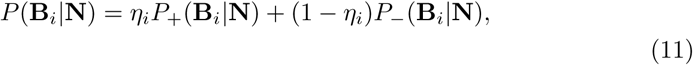

where 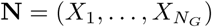 is a vector containing the monomer gene product of all genes, 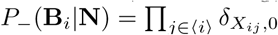, with 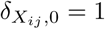 when *X_ij_* = 0, and *P*_+_(**B***_i_*|**N**) is a multinomial PDF, whose generating function is given by:

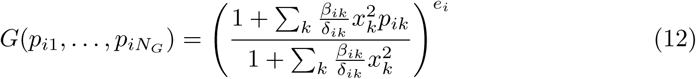

where *e_i_* denotes the number of binding sites at the promoter region of gene *i*, and other parameters are defined in S2 Table (transition rates) and S4 Table (rescaled parameters). The quantity *η_i_* is defined as: *η_i_* = *H*(*Y_i3_* − *Y*_0_) (i.e. gene *i* is epigenetically open if the corresponding level of acetylation, *Y_i3_*, exceeds the threshold *Y*_0_).

Eq. (10) describes the inner dynamics of the fast components (enzymes and enzyme-substrate complexes) of the ER system for each gene *i*. The resulting stochastic dynamics describes how the enzymes switch between their free state and their complex state at constant rates (since the number of the different substrates is constant under the hypothesis of time scale separation). Since the number of enzymes is conserved, the (quasi-)steady distribution of the number of enzymes of each type in complex form is a binomial (see Section 3.4 of the S1 File for a detailed derivation of this result). The corresponding generating functions are given by:

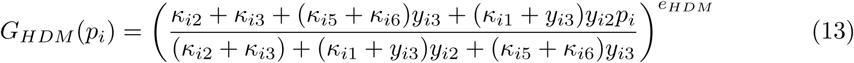

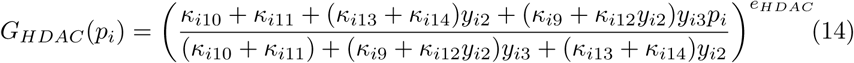

where *i* = 1,…, *N_G_* and *κ_ij_* are defined in S4 Table. The number of free HDM and HDAC molecules is then obtained from the conservation equations *Y_i4_* = *e_HDM_* − *Y_i5_* and *Y_i6_* = *e_HDAC_* − *Y_i7_*.

#### Outer solution

The outer solution, corresponding to the dynamical evolution of the slow variables, is obtained by sampling the fast variables, whose values are needed to compute the reaction rates for the slow variables, from their QSSA PDFs (see Eqs. (12)-(14)):

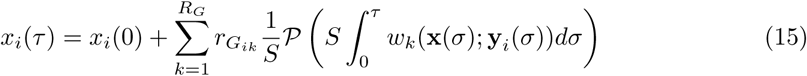

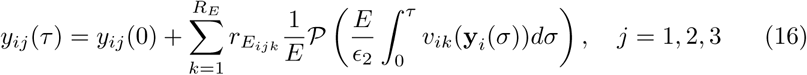

The QSSA PDFs of the fast variables are conditioned by the current value of the slow variables. We complete our asymptotic analysis by looking at the large *S* behaviour of the slow GRN variables (see Eq. (5)). We resort to a law of large numbers enunciated and proved by Kurtz which states that *S*^−1^*P*(*S_u_*) → *u* when *S* ≫ 1 [29-31,51]. We can apply this result straightforwardly to Eq. (15), which eventually leads to the asymptotic reduction of the full ER-GRN system:

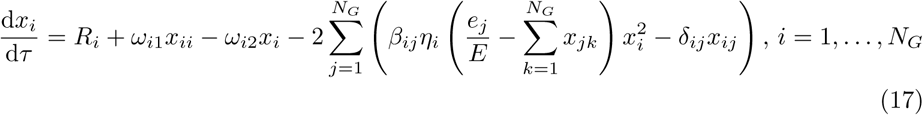

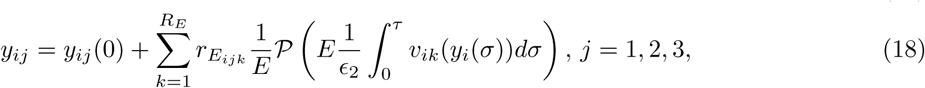

The resulting dynamics consists on a hybrid system where the dynamics of the TF monomers, *x_i_*(*τ*), Eq. (17), is described in terms of a piece-wise deterministic Markov process [52,53], i.e. by a system of ODEs perturbed at discrete times by two random processes, one corresponding to stochastic ER (Eq. (18)) and the other to TF dimers binding to the promoter regions. The latter are sampled from their QSSA PDFs, Eq. (11). The stochastic dynamics of the slow ER variables, Eq. (18) is in turn coupled to the random variation of the associated fast variables (ER enzymes, HDM and HDAC, and complexes). The number of complexes, *Y_i5_* and *Y_i7_*, are sampled from their QSSA PDFs, Eqs. (13) and (14). The corresponding numerical method used to simulate such system is described in detail in S1 Appendix.

### Transitions between ER states: minimum action path approach

Noise-induced transitions are essential to understand ER dynamics and their effect on cell-fate determination [23]. Throughout the bistable regime, sufficiently large fluctuations in the stochastic ER system will induce switching between the open and silenced states. The rate at which such transitions occur can be described using reaction-rate theory [46] and large deviation theory [47], which show that the waiting time between transitions is exponentially distributed. The average switching time, *τ_s_*, increases exponentially with system size, which in this case is given by the scale of ER substrates, *Y* [47,48,55,56]:

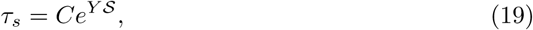

where *C* is a constant and *S* is the action of the stochastic switch. Eq. (19) is derived from considering the probability distribution of the so-called *fluctuation paths*, *φ*(*τ*), which connect the mean-field steady states in a time *τ*. According to large deviation theory [47,48], we have 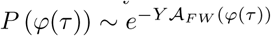, which implies that the probability of observing paths different from the optimal, i.e. the path *φ*_*_ that minimises the action, is exponentially supressed as system size, *Y*, increases. This means that, for large enough system size, the behaviour of the system regarding large fluctuations is characterised by the optimal path, which is such that:

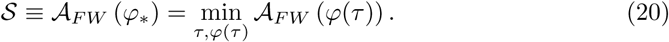

An explicit form of the functional *A_FW_* (*φ*(*τ*)) can be given if the dynamics is given by the corresponding chemical Langevin equation [57]:

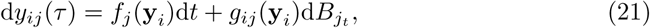

where 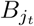 denotes a Wiener process, and the mean-field drift, *f_j_* (y*_i_*), is 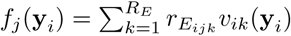, and the noise matrix, *g_ij_*(y*_i_*), 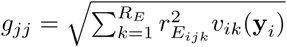, *g_ij_* = 0 if *i* ≠ *j*. The rescaled variables 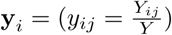, *j* = 1,…, 7, and the rescaled rates *υ_ik_*(y*_i_*) are defined in *Multi-scale analysis and model reduction*. In this case, the action functional *A_FW_*(*φ*(*τ*)) is the Freidlin-Wentzel (FW) functional:

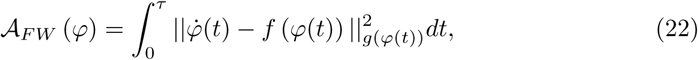

The norm 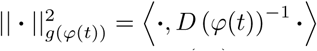, where *D* (*φ*(*t*)) = *g* (*φ*(*t*)) *g* (*φ*(*t*))*^T^* is the diffusion tensor. Using Eq. (22), the optimal value of the action, *S*, can be found by numerical minimisation, which provides both the optimal or minimum action path (MAP) and the rate at which the ER system switch state driven by intrinsic noise. Details regarding implementation of the action-optimisation algorithm are given in Section S.5 of the S1 File. A complete description of *τ_s_* requires to estimate the pre-factor *C*, which is not provided by the FW theory, but can be easily estimated using stochastic simulation.

### ER-systems ensemble generation and analysis

Folguera-Blasco et al. [18] have proposed to analyse an ensemble of ER systems in order to study the robustness of the different ER scenarios under heterogeneous conditions regarding the availability of co-factors associated with the activity of ER enzymes, which we take into account by considering variations (variability) in the kinetic constants *c_ij_* (see S1 Table and S3 Table). Such an ensemble is generated using approximate Bayesian computation (ABC) [58,59], whereby we generate an ensemble of parameter sets *θ_i_* = (*c_ij_*, *i* = 1,…, *N_G_*, *j* = 1,…, 16) compatible with simulated data for the epigenetic regulation systems.

Our approach follows closely that of [18], to which we refer the readers for a detailed presentation of the implementation. To summarise, we start by generating synthetic (simulated) data (denoted as “raw data” in Fig. 3) regarding the ER system of a genetic network epigenetically poised for differentiation, i.e. open differentiation-promoting genes and silenced pluripotency-promoting genes (see the example shown in Fig. 3). This simulated data will play the role of the experimental data, *x*_0_, to which we wish to fit our model. The data set consists of realisations and time points per realisation for each of the *N_G_* epigene regulatory systems. For each time point, *t_i_*, we consider two summary statistics: the mean over realisations, 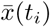, and the associated standard deviation, σ(*t_i_*). We then run the ABC rejection sampler method until we reach an ensemble of parameter sets which fit the simulated data, *x*_0_, within the prescribed tolerances for the mean and standard deviation. In our particular case *N_G_* = 2, i.e. we have a parameter set ensemble of the ER system for the pluripotency-promoting gene and another parameter set ensemble for the gene promoting differentiation. Fig. 3(a) & (b) show results comparing the reference (raw simulated) data to a subensemble average consisting of the sets that best fit the data, for the differentiation- and pluripotency-promoting genes, respectively.

**Fig 3.**
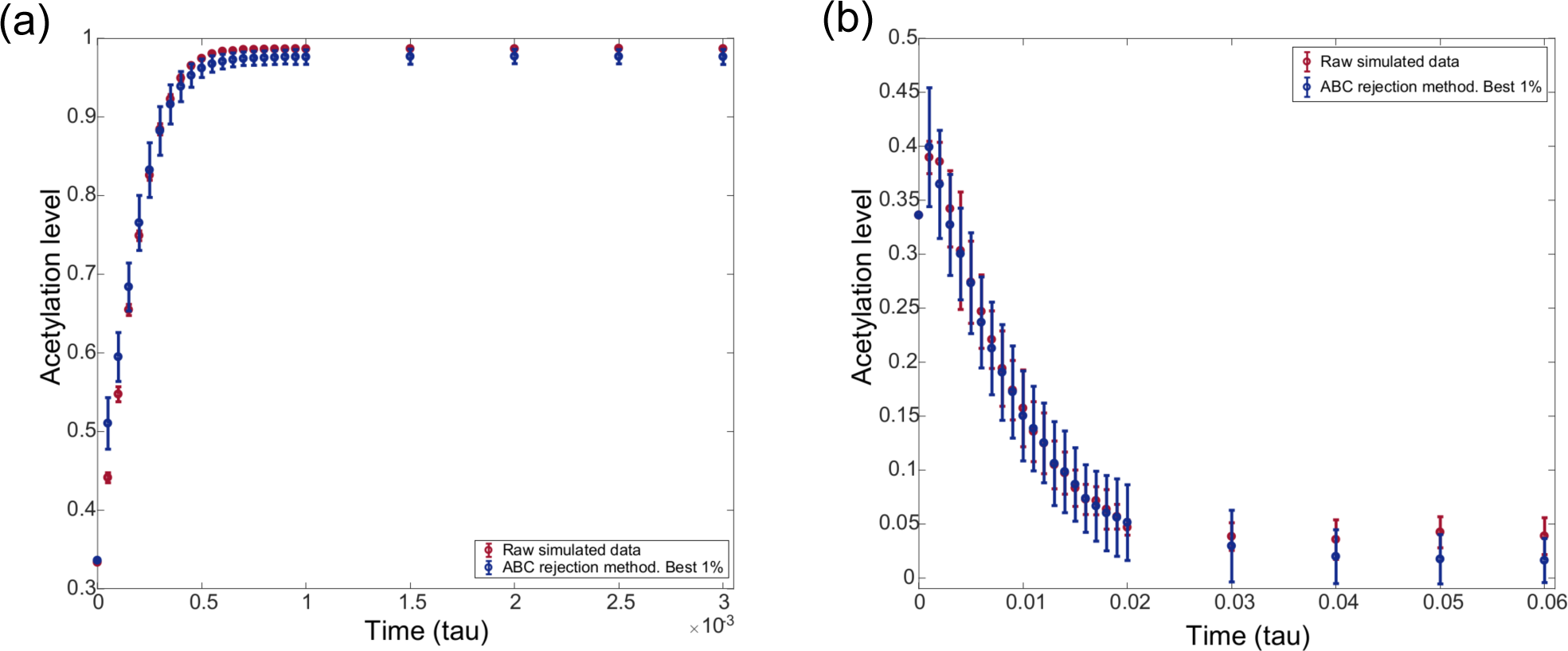
Comparison between raw simulated data (red) with best fitted data resulting from the parametric sensitivity analysis of the epigenetic regulatory system (blue). (a) Evolution to the open state of the differentiation-regulating gene. (b) Evolution to the silenced state of the pluripotency-regulating gene. Raw simulated data is generated by using the SSA on the model defined by the rates shown in S3 Table with parameter values given in Tables S.5 and S.6 in S1 File, for (a) and (b), respectively. Resulting fitted data correspond to the ABC parameter sets that best fit the raw data.

The above procedure provides us with an ensemble of parameter sets that are compatible with our raw data, i.e. such that they fit the data within the prescribed tolerances. The heterogeneity associated with the variability within this ensemble has a clear biological interpretation. The rates *c_ij_* are associated with the activity of the different enzymes that carry out the epigenetic-regulatory modifications (HDMs, HDACs, as well as, histone methylases (HMs) and histone acetylases (HACs)), so that variation in these parameters can be traced back to heterogeneity in the availability of cofactors, many of them of metabolic origin such as NAD_+_, which are necessary for these enzymes to perform their function [18].

The generated kinetic rate constants are dimensionless, i.e. they are relative to a given rate scale [18]. Such a feature implies that there is an undetermined time scale in our system. This additional degree of freedom can be used to fit our model of epigenetic (de-)activation to particular data. Since the global time scales associated with different ER regulation systems may differ among them, our model has the capability of reproducing different systems characterised by different time scales as previously shown by Bintu et al. [14].

## Results

In order to focus our discussion, we illustrate the application of our model formulation on the case we want to study: a gene regulatory circuit with two genes, one whose product promotes differentiation and another one whose protein induces pluripotency. These two genes are further assumed to interact through mutual competitive inhibition (see Fig. 2(a)). Although such a system may appear to be too simplistic to describe realistic situations, there is evidence that mutual inhibition between two key transcription factors controls binary cell fate decisions in a number of situations [60,61]. Our results are straightforward to generalise to more complex situations.

We proceed to analyse how ER sculpts the epigenetic landscape over the substrate of the phase space given by the model of the gene regulatory network. The latter provides the system with a variety of cell fates, corresponding to the stable steady states of the dynamical system underpinning the model of gene regulatory network [62]. The transitions between such cellular states, both deterministic and stochastic, depend upon the ability of the cell regulatory systems to elevate or lower the barriers between them. Epigenetic regulation is one of such mechanisms. Here, we examine how ER is affected by ensemble variability associated with variations in the availability of the necessary co-factors on which histone modifying enzymes (HMEs) depend to carry out their function. In particular, we will show that such variability is enough to produce a variety of behaviours, in particular differentiation-primed and stem-locked states.

### The GRN model exhibits a complex phase space, including an undecided regulatory state

We start our analysis by studying the phase space of the dynamical system underlying our model of gene regulation, schematically illustrated in Fig. 2(a). Using the methodology described in detail in Section S.3 of the S1 File, we have derived the (quasi-steady state approximation) equations for the optimal path theory of the stochastic model of the mutually inhibitory two-gene system [13]. Such equations describe the most likely relaxation trajectories towards their steady states [55,56], under conditions of time scale separations described in detail in Section S.3. For the two GRN, they read as

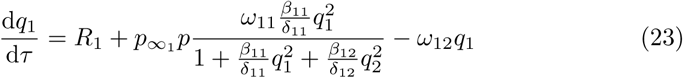

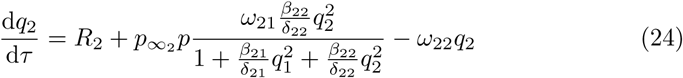

where *q*_1_ and *q*_2_ are the variables (generalised coordinates) associated with the number of molecules of proteins, *X*_1_ and *X*_2_. The re-scaled variables, *q_i_* and *q_j_*, and the re-scaled parameters, *ω_ij_*, *β_ij_*, and *δ_ij_*, are defined in S4 Table (see also Section S.3 in the S1 File).

The multiscale analysis carried out in Section S.3 shows that the parameters *p*, 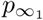 and 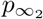 are such that 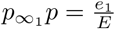 and 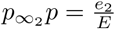, where *e*_1_ and *e*_2_ are the number of sites in the promoter regions of our two genes exposed to and available for binding by TFs, which implies that 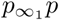 and 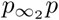 can be directly related to ER: 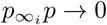, *i* = 1, 2, corresponds to an epigenetically silenced gene, whereas 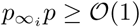 is associated with an epigenetically open gene. In this section, we study the phase space of the system when both 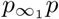 and 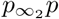 are varied. This allows us to understand how the behaviour of the GRN changes when its components are subject to ER. Our results are shown in Fig. 4.

**Fig 4.**
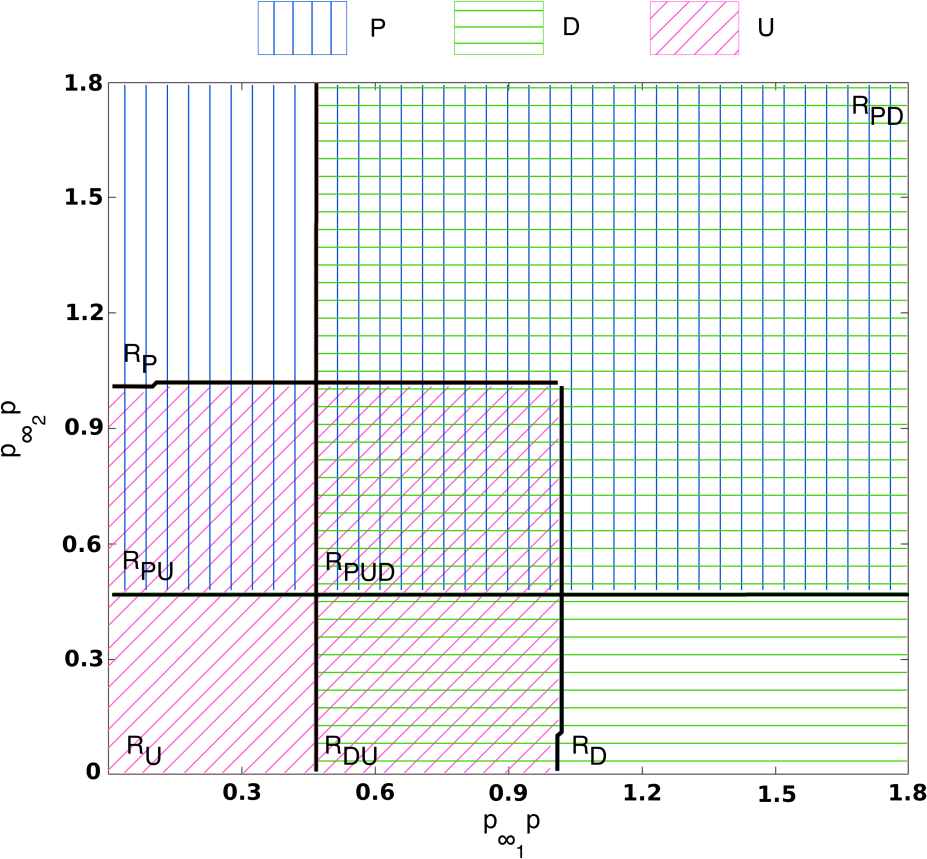
Phase diagram of the two-gene system, Eqs. (23)-(24). Vertical blue (horizontal green) hatching denotes regions where the pluripotency (differentiated) state is stable. Diagonal pink hatching denotes regions where the undecided state is stable. Regions of the phase diagram where different hatchings overlap correspond to regions of bistability or tristability. In the labels in the plot, *P* stands for pluripotency, *D* stands for differentiation and *U* for undecided. This phase diagram was obtained using the methodology formulated in [54]. Parameters values: *ω*_11_ = *ω*_21_ = 4.0. Other parameter values as per Table S.12 in Section S.7 of the S1 File.

The system described by Eqs. (23)-(24) exhibits three types of biologically relevant steady states, namely, the *pluripotency* steady state (PSS), the *differentitation* steady state (DSS), and the *undecided* steady state (USS). Different combinations of these states can be stable or unstable depending on the parameter values (see Fig. 4). The PSS (DSS) corresponds to a steady state with *q*_1_ ≪ 1 and *q*_2_ = O(1) (*q*_1_ = O(1) and *q*_2_ ≫ 1) and the USS is associated with a state such that both *q*_1_ ≪ 1 and *q*_2_ ≪ 1.

Fig. 4 shows the phase space associated with the dynamical system Eqs. (23) and (24). It shows the behaviour as the parameters 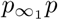 and 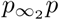 vary. The lines shown in Fig. 4 correspond to the stability boundary of the different regimes. At such boundaries, saddle-node bifurcations occur, as illustrated in the example shown in Fig. S.2 (Section S.4 of the S1 File). Fig. 4 reveals a complex phase space with seven different phases. We denote by *ℛ_P_* (*ℛ_D_*) the region of the phase space where only the pluripotency (differentiation) steady state is stable. Similarly, *ℛ_U_* corresponds to those parameter values such that only the undecided steady state is stable. Furthermore, there are three bistable phases: one in which the PSS and the DSS coexist, *ℛ_PD_*, a second one where the PSS coexists with the USS, *ℛ_PU_*, and the third one where the DSS and USS coexist, *ℛ_DU_*. Finally, a region exists where stable PSS, stable DSS, and stable USS, *ℛ_PUD_*, coexist. Fig. S.5 of the S1 File shows examples of trajectories illustrating the dynamics described by Eqs. (23)-(24) for different values of the pair 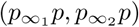 corresponding to the different regions shown in Fig. 4. In particular, we show how the long term behaviour of different initial conditions differ as 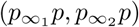 varies, so that different cell fates (co)exist associated to different levels of TF accessibility.

**Fig 5.**
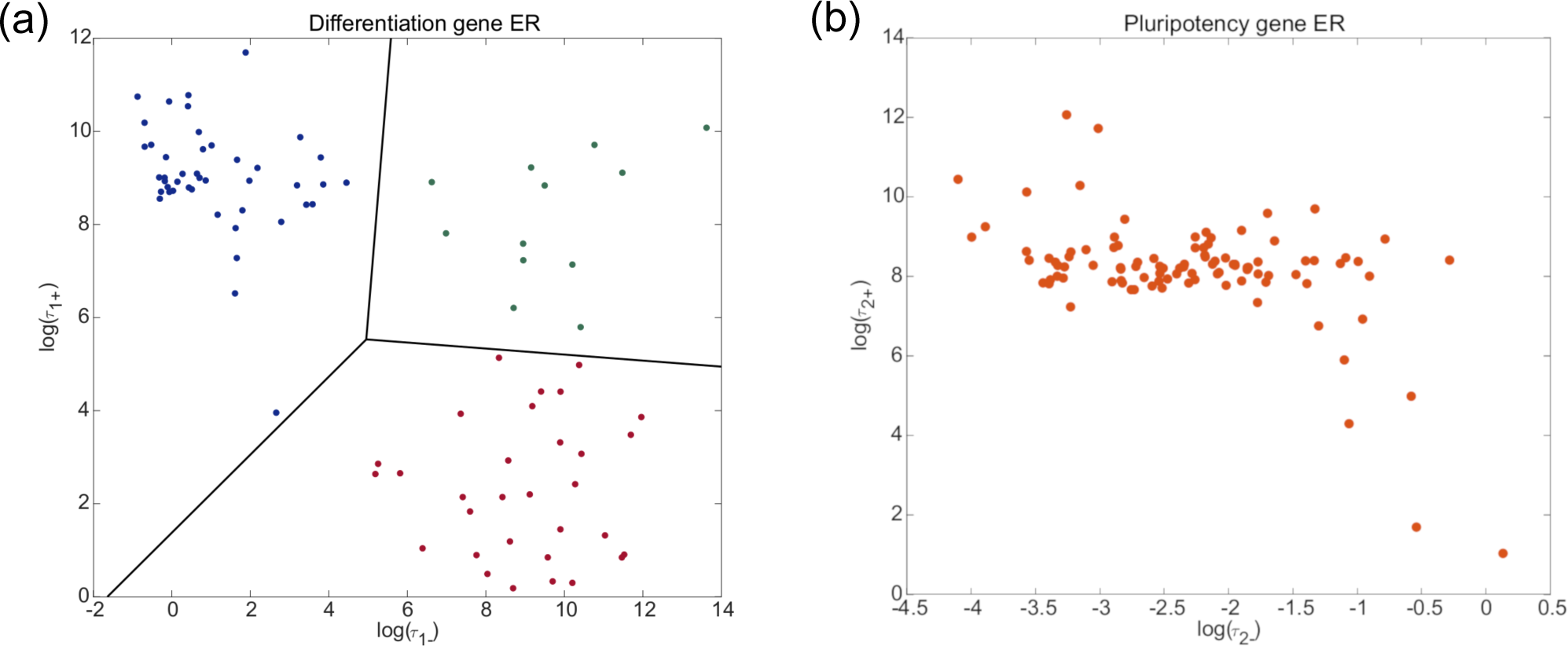
Scatter plots showing heterogeneity in the behaviour of bistable differentiation ER systems (DERSs) and pluripotency ER systems (PERSs). The vertical axis corresponds to the average opening time and the horizontal axis, to the average closing time. Each dot in plot (a) represents a DERS within the ensemble (see Section *ER-systems ensemble generation and analysis*). Different colours and black lines show the three clusters resulting from a *k*-means analysis discussed in Sections *Co-factor heterogeneity gives rise to both pluripotency-locked and differentiation-primed states* and *Analysis of ensemble heterogeneity*. Dots in plot (b) represent PERSs within the ensemble defined in Section *ER-systems ensemble generation and analysis.*

### Co-factor heterogeneity gives rise to both pluripotency-locked and differentiation-primed states

In the previous section, we have analysed the dynamical landscape provided by the dynamical system describing the GRN. We now proceed to study the effect of ER on the robustness of the different phases shown in Fig. 4 (see also Fig. S.5, Section S.8 of the S1 File). We here put forward that ER is essential to the robustness of such phases and, consequently, to the stability of the associated cell fates, since transitions in bistable ER systems can induce (or facilitate) transitions between the GRN phases. Such transitions are associated with differentiation and reprogramming of cell fates. This phenomenon, so-called *epigenetic plasticity*, has been recently proposed as a major driver for disrupting cell-fate regulatory mechanisms in cancer and aging [23]. We further focus on the role of heterogeneity within the ensemble described in Section *Materials and methods* (see also [18]).

In order to characterise robustness of the different ER systems within the ensemble, we have focused on the analysis of the average transition times between the *open* and *closed* ER states. We define 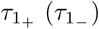 as the average transition time for a differentiation ER system-DERS-to switch from closed to open (open to closed). Similarly, the quantities 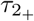 and 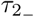 are analogously defined for the pluripotency ER systems-PERSs. The results are shown in Fig. 5(a) and (b), where we present scatter plots of the average transition times within the ensemble of DERSs (Fig. 5(a)) and PERSs (Fig. 5(b)). These figures show scatter plots where each point represents an ER system (i.e. a given parameter set) within our ensemble. The vertical and horizontal axes show the average switching time from closed-to-open and open-to-closed, respectively. We observe that the heterogeneity exhibited by the differentiation ER systems is greater than the one corresponding to the pluripotency ER systems. In particular, the dispersion in 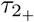 is much smaller than in 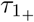. Regarding 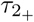, most of the pluripotency ER systems are concentrated around a narrow band. By contrast, the differentiation ER systems show large degrees of heterogeneity in both 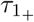 and 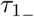.

Heterogeneity in the differentiation ER systems exhibits an interesting pattern, whereby such systems organise themselves in three clusters obtained through *k*-means clustering, shown as blue, green and red dots in Fig. 5(a). DERSs within the *blue* cluster are charaterised by long closed-to-open waiting times and short open-to-closed waiting times. DERSs belonging to the *red* cluster are the specular image of those within the blue cluster, i.e. they have short closed-to-open waiting times and long open-to-closed waiting times. Finally, DERSs in the *green* cluster are characterised by large values of both 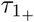 and 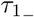.

Insight into the stochastic dynamics, particularly regarding heterogeneity of the robustness of the open and silenced ER states to intrinsic noise, can be gained by analysing the corresponding optimal escape paths. Four examples of such paths, computed according to the MAP theory (see Section *Transitions between ER states: minimum action path approach*), for two DERSs (DERS1 and DERS2) and two PERSs (PERS1 and PERS2) are shown in Figs. S.6(a)-(d) of the S1 File. A comparison between the value of the minimum action, *S* (see Eqs.(20)-(22)), for the optimal escape paths corresponding to DERS1 and DERS2 (Figs. S.6(a) and (c)), and for PERS1 and PERS2 (Figs. S.6(b) and (d)) shows a tendency for DERSs to exhibit much more variability (see Table 1). Whilst the action value for PERS1 is about twice the value of PERS2 in Fig. S.6(b), there is an over 8-fold increase when comparing the action values of DERS1 and DERS2 in Fig. S.6(a). Similarly, when comparing the action *S* for the open to closed optimal paths, we observe that the variability associated with the DERSs (Fig. S.6(c)) is also larger than the one in PERSs (Fig. S.6(d)). This property partly explains the difference between Fig. 5(a) and Fig. 5(b) regarding DERS and PERS heterogeneity, respectively. A similar argument can be put forward to help us explain the heterogeneity within the DERS ensemble (Fig. 5(a)). Blue cluster DERSs exhibit optimal closed-to-open paths with larger value of the optimal action than that found in their red cluster counterparts (see Fig. S.6(a), where DERS1 belongs to the red cluster and DERS2 to the blue cluster). This property has the consequence that the closed-to-open waiting time, 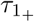, is longer for blue cluster DERSs.

**Fig 6.**
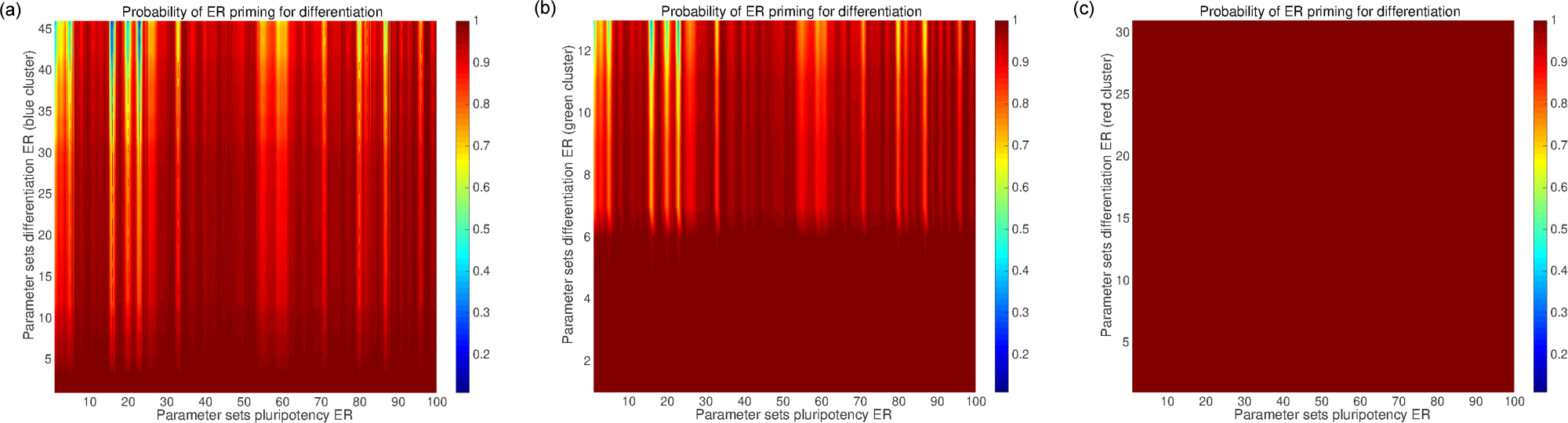
Differentiation probability *Q* within the ensembles of DERSs and PERSs corresponding to the three clusters of Fig. 5(a) (see Section *Co-factor heterogeneity gives rise to both pluripotency-locked and differentiation-primed states*). For all three plots, the horizontal axis runs over the whole ensemble of PERSs. The vertical axis of plots (a), (b), and (c) runs over all the DERSs within the blue cluster, the green cluster, and the red cluster, respectively.

**Table 1.**
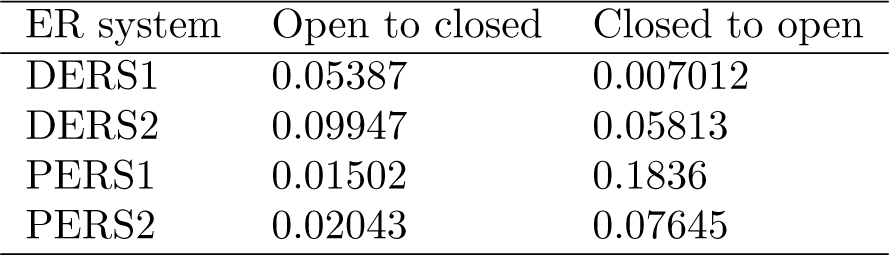
Minimum action values, *S*, corresponding to the optimal escape paths shown in Fig. S.6 of the S1 File (see Section *Transitions between ER states: minimum action path approach* and Section *Co-factor heterogeneity gives rise to both pluripotency-locked and differentiation-primed states* for details). Parameter values are given in Section S.7 of the S1 File.

**Table 2.**
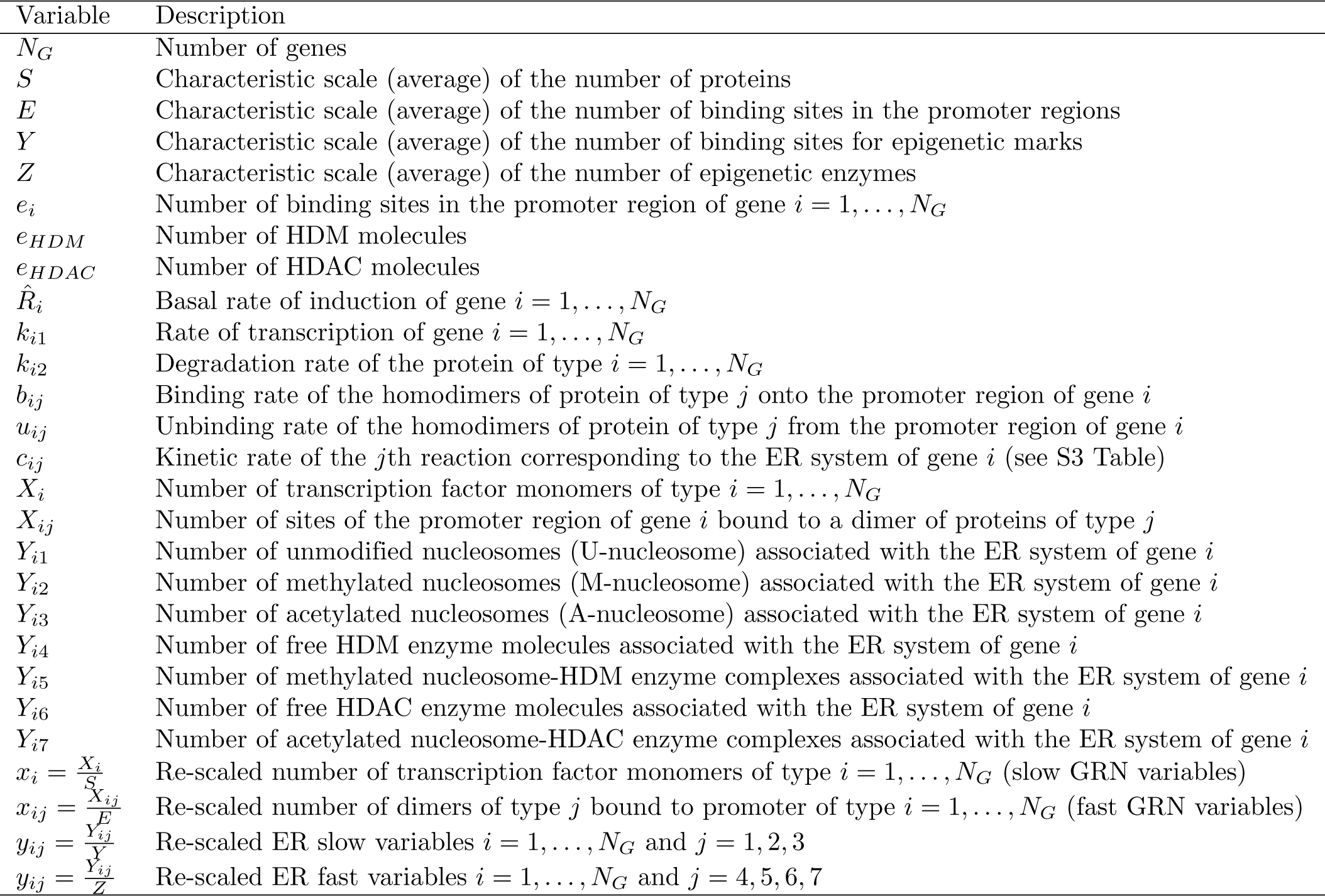
Brief description of the variables and parameters involved in the symmetric model of gene regulatory network with competitive binding inhibition. For simplicity, we will assume that *O*(*E*) = *O*(*Y*)

| Variable | Description |
| --- | --- |
| $N_G$ | Number of genes |
| $S$ | Characteristic scale (average) of the number of proteins |
| $E$ | Characteristic scale (average) of the number of binding sites in the promoter regions |
| $Y$ | Characteristic scale (average) of the number of binding sites for epigenetic marks |
| $Z$ | Characteristic scale (average) of the number of epigenetic enzymes |
| $e_i$ | Number of binding sites in the promoter region of gene $i = 1, \dots, N_G$ |
| $e_{HDM}$ | Number of HDM molecules |
| $e_{HDAC}$ | Number of HDAC molecules |
| $\hat{R}_i$ | Basal rate of induction of gene $i = 1, \dots, N_G$ |
| $k_{i1}$ | Rate of transcription of gene $i = 1, \dots, N_G$ |
| $k_{i2}$ | Degradation rate of the protein of type $i = 1, \dots, N_G$ |
| $b_{ij}$ | Binding rate of the homodimers of protein of type $j$ onto the promoter region of gene $i$ |
| $u_{ij}$ | Unbinding rate of the homodimers of protein of type $j$ from the promoter region of gene $i$ |
| $c_{ij}$ | Kinetic rate of the $j$th reaction corresponding to the ER system of gene $i$ (see S3 Table) |
| $X_i$ | Number of transcription factor monomers of type $i = 1, \dots, N_G$ |
| $X_{ij}$ | Number of sites of the promoter region of gene $i$ bound to a dimer of proteins of type $j$ |
| $Y_{i1}$ | Number of unmodified nucleosomes (U-nucleosome) associated with the ER system of gene $i$ |
| $Y_{i2}$ | Number of methylated nucleosomes (M-nucleosome) associated with the ER system of gene $i$ |
| $Y_{i3}$ | Number of acetylated nucleosomes (A-nucleosome) associated with the ER system of gene $i$ |
| $Y_{i4}$ | Number of free HDM enzyme molecules associated with the ER system of gene $i$ |
| $Y_{i5}$ | Number of methylated nucleosome-HDM enzyme complexes associated with the ER system of gene $i$ |
| $Y_{i6}$ | Number of free HDAC enzyme molecules associated with the ER system of gene $i$ |
| $Y_{i7}$ | Number of acetylated nucleosome-HDAC enzyme complexes associated with the ER system of gene $i$ |
| $x_i = \frac{X_i}{S}$ | Re-scaled number of transcription factor monomers of type $i = 1, \dots, N_G$ (slow GRN variables) |
| $x_{ij} = \frac{X_{ij}}{E}$ | Re-scaled number of dimers of type $j$ bound to promoter of type $i = 1, \dots, N_G$ (fast GRN variables) |
| $y_{ij} = \frac{Y_{ij}}{Y}$ | Re-scaled ER slow variables $i = 1, \dots, N_G$ and $j = 1, 2, 3$ |
| $y_{ij} = \frac{Y_{ij}}{Z}$ | Re-scaled ER fast variables $i = 1, \dots, N_G$ and $j = 4, 5, 6, 7$ |

**Table 3.**
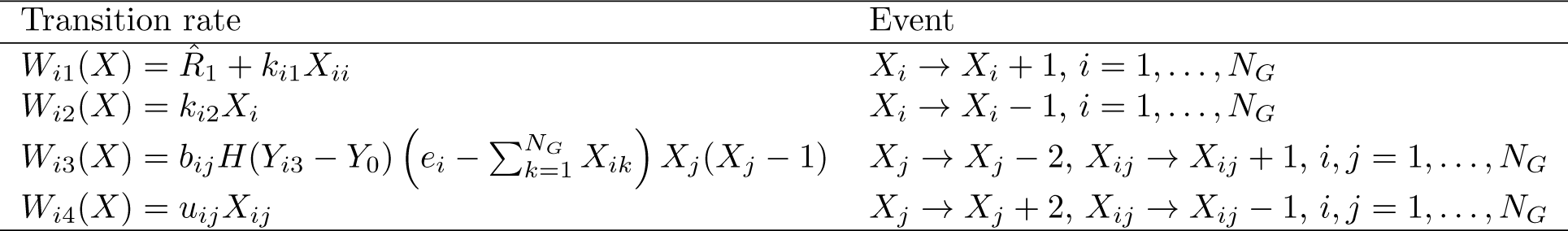
Transition rates associated with the stochastic dynamics of gene regulatory circuit. Note that the rate of binding of homodimers to the promoter region of gene *i* is modulated by the level of acetylation of gene *i*, *Y_i3_*: if *Y_i3_* is above the threshold *Y*_0_, gene *i* is *open*, i.e. the promoter is accessible to homodimers and TFs. By contrast, if the gene’s acetylation levels decay, gene *i* is *silenced* and the promoter is inaccessible to gene-transcription regulatory dimers.

| Transition rate | Event |
| --- | --- |
| $W_{i1}(X) = \hat{R}_1 + k_{i1}X_{ii}$ | $X_i \rightarrow X_i + 1, i = 1, \dots, N_G$ |
| $W_{i2}(X) = k_{i2}X_i$ | $X_i \rightarrow X_i - 1, i = 1, \dots, N_G$ |
| $W_{i3}(X) = b_{ij}H(Y_{i3} - Y_0) \left( e_i - \sum_{k=1}^{N_G} X_{ik} \right) X_j(X_j - 1)$ | $X_j \rightarrow X_j - 2, X_{ij} \rightarrow X_{ij} + 1, i, j = 1, \dots, N_G$ |
| $W_{i4}(X) = u_{ij}X_{ij}$ | $X_j \rightarrow X_j + 2, X_{ij} \rightarrow X_{ij} - 1, i, j = 1, \dots, N_G$ |

**Table 4.** This table shows the transition rates associated with the stochastic dynamics of the epigenetic regulatory system of gene *i*. The random variables *Y_ik_* are defined in S1 Table. The different modification reactions are assumed to be of two types, recruited and unrecruited. Details regarding the assumptions between this distinction as well as a full description of the formulation of the stochastic epigenetic regulation model are given in Section S.1, S1 File. We also refer the reader to references [13,18,19].

| Transition rate | Reaction change vector | Event |
| --- | --- | --- |
| $V_{i1} = c_{i1}Y_{i2}Y_{i4}$ | $r_{E_{i1}} = (0, -1, 0, -1, +1, 0, 0)$ | Formation of M-nucleosome-HDM enzyme complex (unrecruited) |
| $V_{i2} = c_{i2}Y_{i5}$ | $r_{E_{i2}} = (0, +1, 0, +1, -1, 0, 0)$ | M-nucleosome-HDM enzyme complex splits (unrecruited) |
| $V_{i3} = c_{i3}Y_{i5}$ | $r_{E_{i3}} = (+1, 0, 0, +1, -1, 0, 0)$ | Demethylation and HDM enzyme release (unrecruited) |
| $V_{i4} = c_{i4}Y_{i2}Y_{i3}Y_{i4}$ | $r_{E_{i4}} = (0, -1, 0, -1, +1, 0, 0)$ | Formation of M-nucleosome-HDM enzyme complex (recruited) |
| $V_{i5} = c_{i5}Y_{i3}Y_{i5}$ | $r_{E_{i5}} = (0, +1, 0, +1, -1, 0, 0)$ | M-nucleosome-HDM enzyme complex splits (recruited) |
| $V_{i6} = c_{i6}Y_{i3}Y_{i5}$ | $r_{E_{i6}} = (+1, 0, 0, +1, -1, 0, 0)$ | Demethylation and HDM enzyme release (recruited) |
| $V_{i7} = c_{i7}Y_{i1}$ | $r_{E_{i7}} = (-1, +1, 0, 0, 0, 0, 0)$ | Methylation (unrecruited) |
| $V_{i8} = c_{i8}Y_{i1}Y_{i2}$ | $r_{E_{i8}} = (-1, +1, 0, 0, 0, 0, 0)$ | Methylation (recruited) |
| $V_{i9} = c_{i9}Y_{i3}Y_{i6}$ | $r_{E_{i9}} = (0, 0, -1, 0, 0, -1, +1)$ | Formation of A-nucleosome-HDAC enzyme complex (unrecruited) |
| $V_{i10} = c_{i10}Y_{i7}$ | $r_{E_{i10}} = (0, 0, +1, 0, 0, +1, -1)$ | A-nucleosome-HDAC enzyme complex splits (unrecruited) |
| $V_{i11} = c_{i11}Y_{i7}$ | $r_{E_{i11}} = (+1, 0, 0, 0, 0, +1, -1)$ | Deacetylation and HDAC enzyme release (unrecruited) |
| $V_{i12} = c_{i12}Y_{i3}Y_{i2}Y_{i6}$ | $r_{E_{i12}} = (0, 0, -1, 0, 0, -1, +1)$ | Formation of A-nucleosome-HDAC enzyme complex (recruited) |
| $V_{i13} = c_{i13}Y_{i7}Y_{i2}$ | $r_{E_{i13}} = (0, 0, +1, 0, 0, +1, -1)$ | A-nucleosome-HDAC enzyme complex splits (recruited) |
| $V_{i14} = c_{i14}Y_{i7}Y_{i2}$ | $r_{E_{i14}} = (+1, 0, 0, 0, 0, +1, -1)$ | Deacetylation and HDAC enzyme release (recruited) |
| $V_{i15} = c_{i15}Y_{i1}$ | $r_{E_{i15}} = (-1, 0, +1, 0, 0, 0, 0)$ | Acetylation (unrecruited) |
| $V_{i16} = c_{i16}Y_{i1}Y_{i3}$ | $r_{E_{i16}} = (-1, 0, +1, 0, 0, 0, 0)$ | Acetylation (recruited) |

**Table 5.**
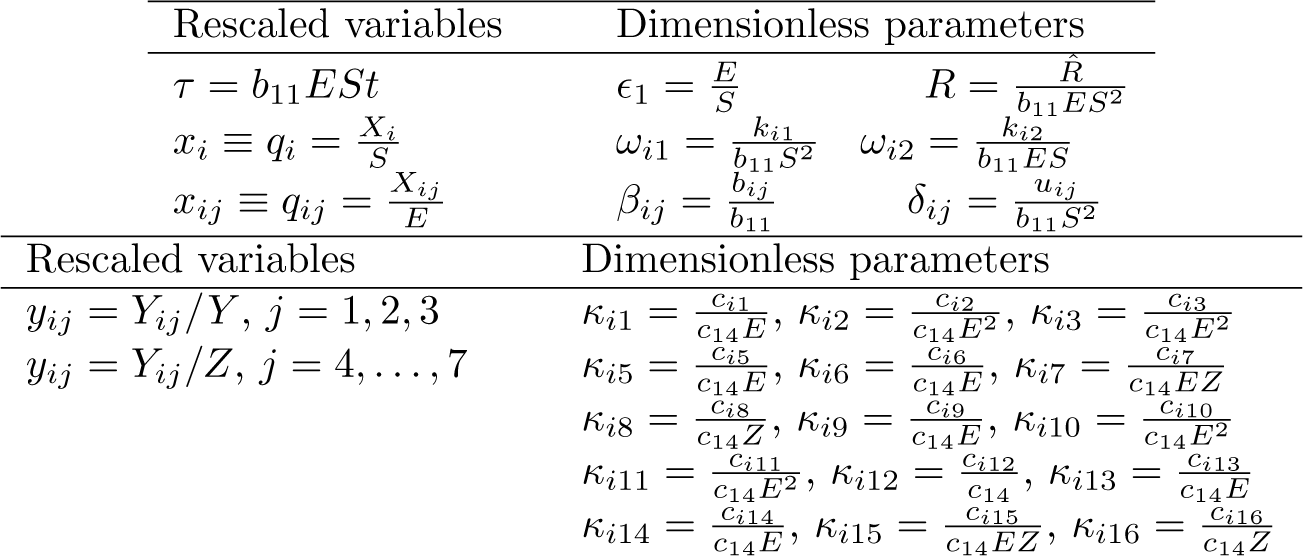
Re-scaled GRN and ER parameters (see Section *Multi-scale analysis and model reduction* and S1 File for details)

To quantify the effects of bistable ER on the landscape related to the gene regulatory system (see Fig. 4), we proceed to estimate the probability, *Q*, that the combined activity of each pair of DERS and PERS within our ensemble produces a global epigenetic regulatory state compatible with differentiation. DERS-PERS pairs with high values of *Q* are associated with *differentiation-primed* states. By contrast, those DERS-PERS combinations with low *Q* are identified with *pluripotency-locked* states.

We proceed forward with this programme by recalling that escape times from a stable attractor in a stochastic multi-stable system are exponentially distributed [47,48]. This implies that the PDFs for the escape times for both DERSs and PERSs are fully determined by the corresponding values of 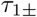 and 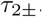. We also assume that, for a given ER-GRN system, the DERS and the PERS evolve independently of each other.

We consider the PDF of the waiting time associated with a scenario of full reprogramming of the epigenetic landscape, *τ_P_*. Such a scenario assumes that the system is initially in a pluripotency-locked ER state where the DERS is closed and the PERS is open, which we denote as *D*_−_*P*_+_. For the system to make its transit into the differentiation-primed state *D*_+_*P*_−_, corresponding to open DERS and closed PERS, there are two possible routes: *D*_−_*P*_+_→ *D*_−_*P*_−_ → *D*_+_*P*_−_ (route 1) and *D*_−_*P*_+_ → *D*_+_*P*_+_ → *D*_+_*P*_−_ (route 2). Simultaneous switch of both ER systems is considered highly unlikely and therefore ignored. The PDF of the waiting time of the transition *D*_−_*P*_+_ → *D*_+_*P*_−_ denoted by *P*_+−,−+_ (*τ_P_*), is given by:

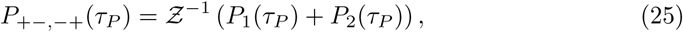

where

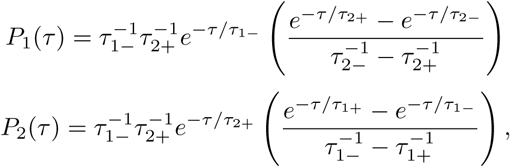

and

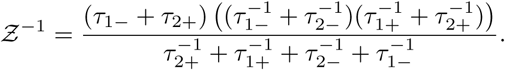

*P*_1_(*τ_p_*) and *P*_2_(*τ_p_*) are the probabilities related to each of the landscape reprogramming routes. The probability that the ER landscape has undergone reprogramming from pluripotency-locked into differentiation-primed state within the time interval (0, *τ_P_*], *Q*, is thus given by:

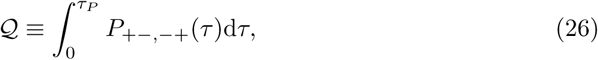

where in our case, *τ_P_* has been taken as the mean time of *τ*_1+_, that is, the mean time for the differentiation ER systems (DERSs) to switch from the closed to the open state, which is needed for a cell to differentiate. Furthermore, *τ*_1+_ exhibits a larger range of variability than the time for the pluripotency ER systems to switch from its open to its closed state, which is also a necessary condition for differentiation to happen.

We investigate the DERSs belonging to the different clusters of Fig. 5(a) regarding their likelihood to produce pluripotency-locked epigenetic landscapes (results shown in Fig. 6). The analysis shows that DERSs within the red cluster (Fig. 6(c)) correspond to differentiation-primed epigenetic landscapes (*Q* = 1) for all the DERS-PERS pairs. By contrast, the blue cluster (Fig. 6(a)) and the green cluster (Fig. 6(b)) contain DERSs associated with both differentiation-primed (large *Q*) and pluripotency-locked (small *Q*) epigenetic landscapes. As discussed in the next section, the latter are more abundant within the blue cluster.

### Analysis of ensemble heterogeneity

We now proceed to analyse the patterns observed in our ensemble of ER systems regarding both the differences between the three clusters observed in the ensemble of DERSs (Fig. 5(a)) and the distinctive features that characterise pluripotency-locked DERS-PERS pairs. In order to do this, we follow the methodology put forward by [18], whereby ensemble statistics (cumulative distribution functions (CDFs)) of the parameters *c_ij_* (see S1 Table) corresponding to the DERSs/PERSs associated with the subensemble of systems exhibiting a particular behaviour are analysed. By comparing such CDFs to either the general population (i.e. whole ensemble) or to different subensembles, we can detect statistically significant biases, which allows us to identify key parameters (and their biases) associated with the behaviour displayed by the focal subensemble.

#### Significant differences within the ensemble of DERSs

We start this analysis by studying the pattern emerging in the ensemble of DERSs, Fig. 5(a). As discussed in the previous section, DERSs organise themselves in three clusters, 566 which exhibit remarkable differences regarding their capability to trigger differentiation-primed epigenetic landscapes (see Figs. 5(a) and 6). Our results are shown in Fig. 7, where we depict the empirical CDFs for the different kinetic parameters of the ER reactions for the differentiation gene, 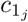 (see S3 Table). We proceed to look for which 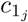 there are statistically significant differences, by comparing the CDF of each cluster with that corresponding to the whole DERS ensemble, and also the CDFs of the clusters among them (see Fig. 7). Each of these two-sample comparison is carried out by means of the Kolmogorov-Smirnov (KS) test. Statistically significant differences were found in the cases we comment below. The *p*-values are reported in Section S.6 of the S1 File.

**Fig 7.**
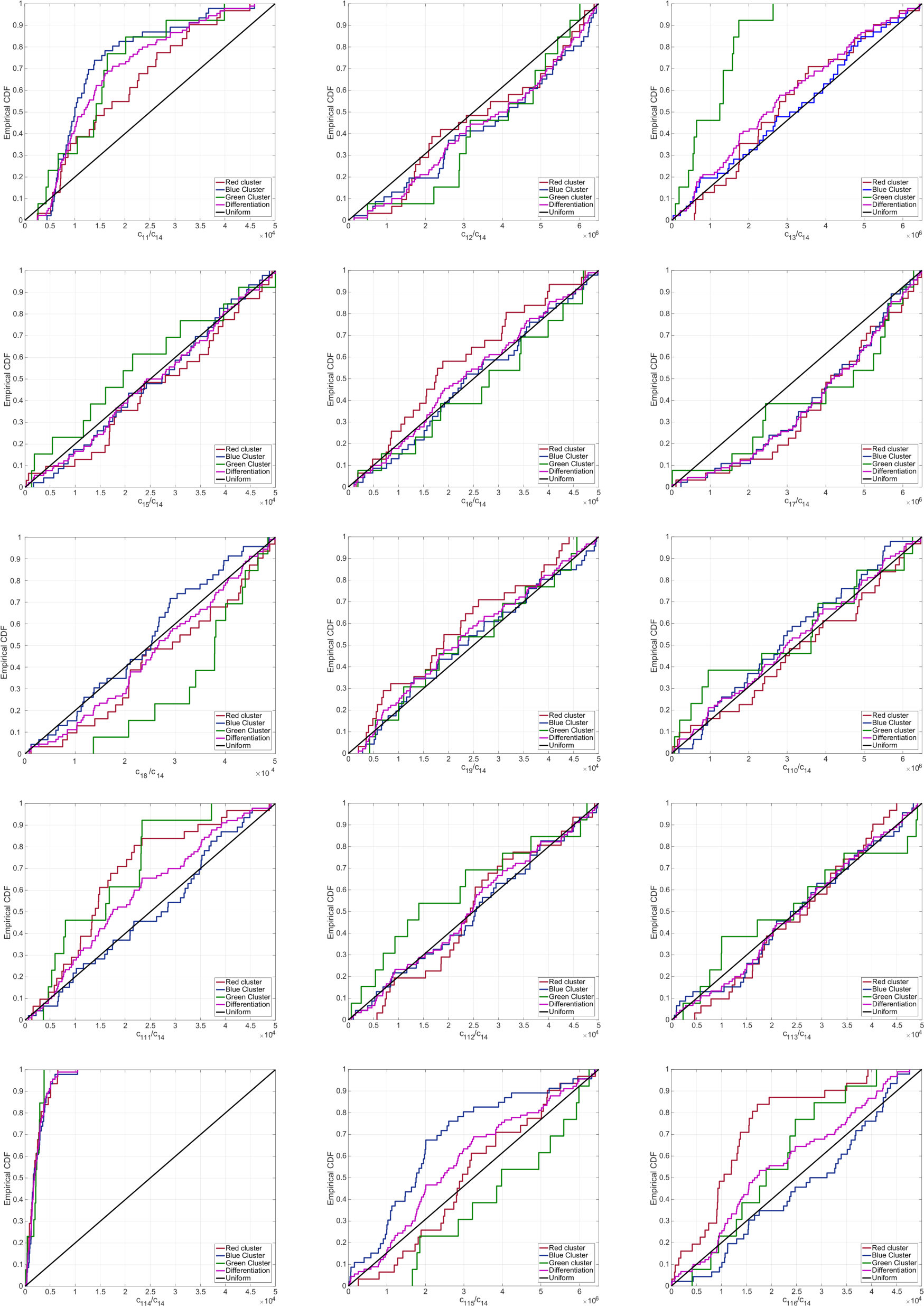
Empirical CDFs for the whole ensemble of DERS parameter sets (magenta lines). This ensemble has been generated according to the methodology explained in Section *ER-systems ensemble generation and analysis* (see also [18]). We also show the partial empirical CDFs corresponding to each of the clusters from Fig. 5(a) (red, green, and blue lines). We analyse a total of DERS parameter sets. The red cluster includes sets, the green cluster contains sets, and the blue cluster has sets. For reference, we also show the CDF for a uniform distribution (black line).

##### Red cluster versus blue cluster

As discussed in the previous section, the differences between DERSs within the blue and red clusters are essential to ascertain the main features that distinguish differentiation-primed and pluripotency-locked systems. The bias detected within the red (blue) cluster in the corresponding CDFs (see Fig. 7) is towards bigger (smaller) values for 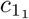 (unrecruited demethylation) and 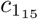 (unrecruited acetylation) and towards smaller (larger) values for 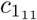 (unrecruited deacetylation) and 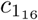 (recruited acetylation). The behaviour of 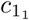, 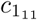, and 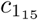 is straightforward to interpret. The trends observed in the data are consistent with the DERSs within red cluster being more prone to differentiation-primed ER landscapes, as they promote removal of negative marks and addition of positive marks.

##### Red cluster versus green cluster

In this case, the bias detected within the red (green) cluster in the corresponding CDFs (see Fig. 7) is to larger (smaller) values for 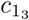 (unrecruited demethylation) and to smaller (bigger) values for 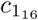 (recruited acetylation). The tendency in the data corresponding to 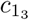 is compatible with the features of the red cluster DERSs, as it involves an increase in the removal of negative marks.

##### Blue cluster versus green cluster

Fig. 7 shows that DERSs within the green cluster have smaller values of 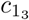 (unrecruited demethylation) and larger values of 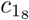 (recruited methylation) than their blue cluster counterparts. Both of such effects stimulate addition of negative marks. However, DERSs in the green cluster also exhibit lower 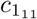 (unrecruited deacetylation) and bigger 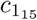 (unrecruited acetylation), which both encourage addition of positive marks. This can explain why the green cluster DERSs exhibit both long 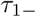 and 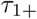 (see Fig. 5(a)).

#### Significant differences between differentiation-primed and pluripotency-locked ER landscapes

The quantity *Q* allows us to classify each pair DERS-PERS drawn from our ensemble regarding their degree of resilience to switch into a state prone to differentiation. If *Q* is larger than a threshold value *T*, the corresponding DERS-PERS pair is categorised as differentiation-primed. By contrast, when *Q* < *T*, the DERS-PERS pair is classified as pluripotency-locked.

We first proceed to compare within the whole population (without discriminating between clusters) those DERSs such that *Q* ≥ *T* (differentiation-primed ER landscapes) against those with *Q* < *T* (pluripotency-locked ER landscapes). We take *T* = 0.7. The results are shown in Fig. S.7 of the S1 File in Section S.8. The CDFs of the parameters 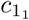 (unrecruited demethylation), 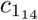 (recruited deacetylation), and 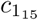 (unrecruited acetylation) are biased towards higher values for the subensemble associated with differentiation-primed ER landscapes (*Q* ≥ *T*). The requirement for *Q* to be *Q* ≥ *T* biases the CDF of 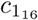 (recruited acetylation) towards lower values than in the general population. The interpretation of the results regarding 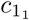 and 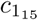 is clear, since they encourage the removal of negative marks and the addition of positive marks and thus promote expression of the differentiation gene. The CDFs of 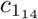 and 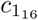 corresponding to differentiation-primed ER landscapes are virtually identical to the CDFs associated with the general population (see Fig. S.7, S1 File). These features are therefore inherent in bistable behaviour (see Section *General description of the stochastic model of an epigenetically-regulated gene network*), rather than being specific to differentiation-primed DERSs.

If we now restrict our analysis to those DERSs within the blue cluster (see Fig. 8), we observe that the parameters whose CDFs differ significantly when splitted into differentiation-primed and pluripotency-locked are 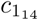 (unrecruited demethylation) and c_ll4_ (recruited deacetylation). As in the analysis in the whole ensemble, only the result regarding 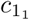 is relevant for the analysis of the features yielding differentiation-primed ER landscapes.

**Fig 8.**
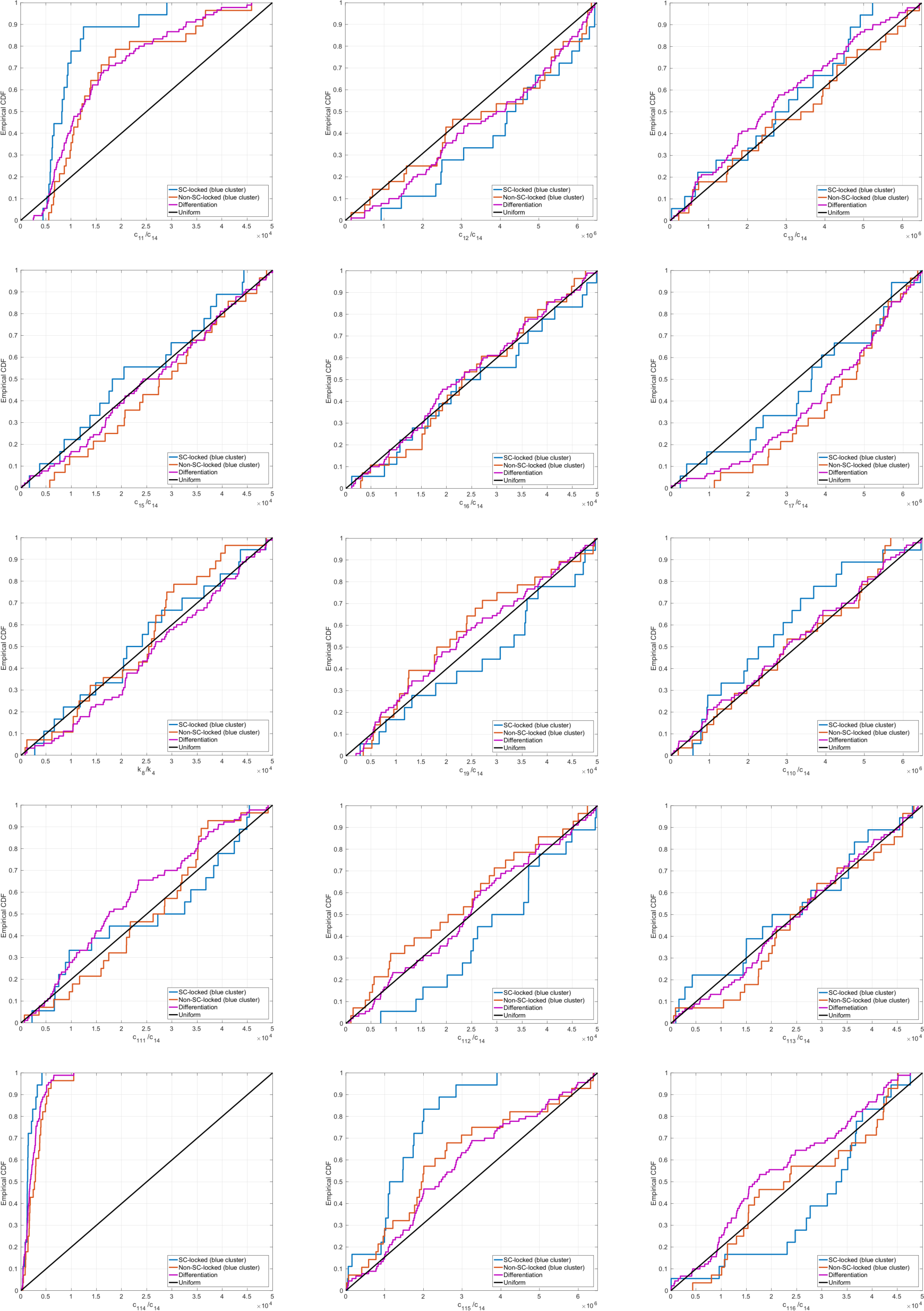
Empirical CDFs for the DERS parameter sets within the blue cluster. This ensemble has been generated according to the methodology explained in Section *ER-systems ensemble generation and analysis* (see also [18]). The DERSs within the blue cluster have been divided into two subsets: those such that *Q* < *T* (SC-locked, blue lines) and those such that *Q* ≥ *T* (non-SC-locked, orange lines), with *T* = 0.7. For comparison, we plot the CDFs of the whole DERS ensemble (magenta lines), and, for guidance the CDF corresponding to a uniformly distributed random variable (black lines).

Regarding the PERSs, the results are less compelling. The results are shown in Fig. S.8 of the S1 File. Our analysis shows that significative differences can be found between the empirical distributions of three parameter values: 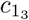 (unrecruited demethylation), 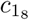 (recruited methylation), and 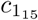 (unrecruited acetylation). PERSs such that *Q* ≥ *T* exhibit larger values of all three parameters.

#### Ensemble-based strategies for unlocking resilient pluripotency

The results of the previous sections suggest a number of strategies to unlock resilient pluripotency states which hinder differentiation. One of our main conclusions is that such states of resilient pluripotency are mostly vinculated to DERS-PERS combinations such that the DERS belongs to either the blue or the green cluster. In view of this, a possible strategy in order to encourage differentitation-primed ER landscape consists on changing a selected combination of parameter values according to a rationale provided by the analysis carried out in the previous two sections. Our results are shown in Fig. 9.

**Fig 9.**
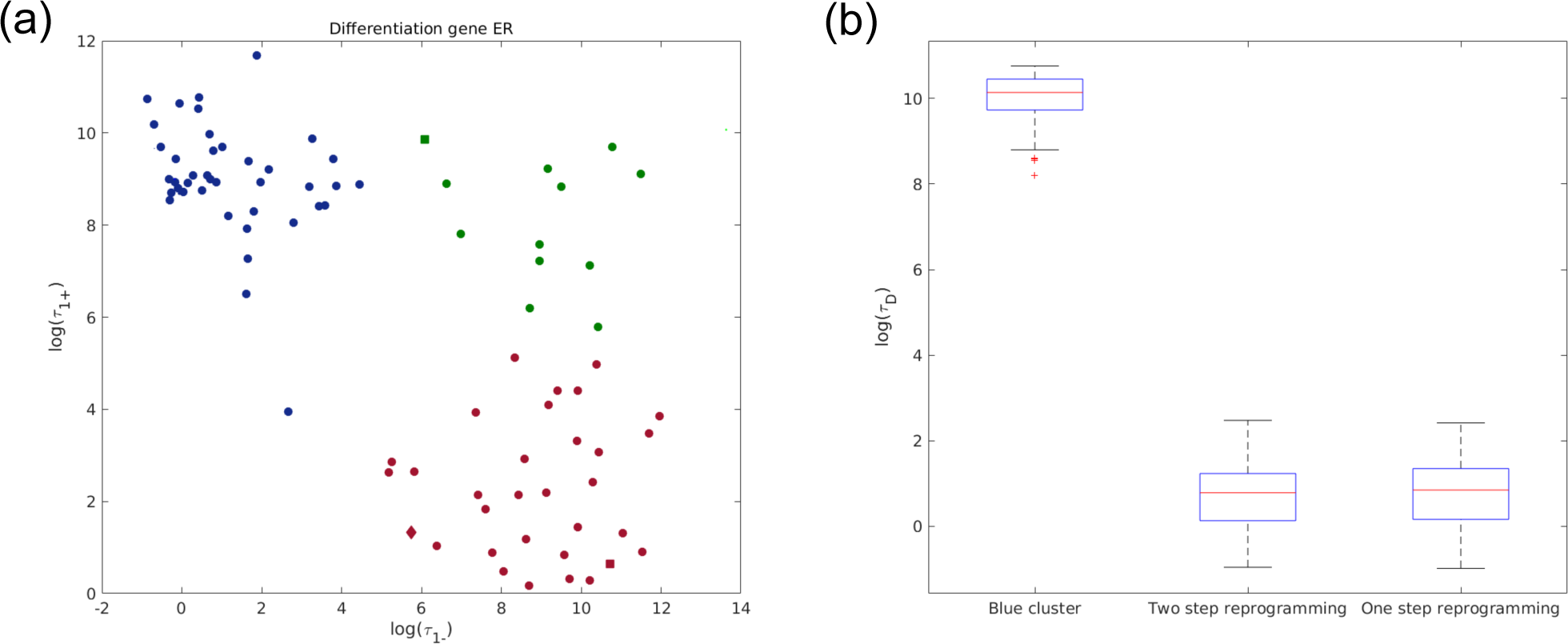
Plots showing the effect of the different reprogramming strategies of blue cluster DERSs, as evaluated in terms of the statistics of the differentiation time (*τ_D_*). (a) Two step reprogramming is illustrated by the green square (first step), which finally becomes the red square (second step). One step reprogramming is depicted as the red diamond (see Section *Ensemble-based strategies for unlocking resilient pluripotency* for details). (b) Comparison of *τ_D_* for the original DERS and the ones resulting from the reprogramming strategies. We consider a base-line scenario where the number of HMEs is exactly equal to average, i.e. *e_HDM_* = *e_HDAC_* = *Z*. We then compare the simulation results obtained for different scenarios regarding the different strategies to the base-line scenario. Parameter values: *Z* = 5 and *Y* = 15. Other parameter values given in Table S.11, Section S.7 of the S1 File.

One possible strategy consists on first transforming a blue cluster DERS into a green cluster one, and then completing the *reprogramming* of the DERS by transforming the resulting set into a red cluster DERS. A candidate strategy involves first changing a parameter whose CDF is significantly different when the blue cluster is compared with the green cluster. The second step is then to change a parameter that exhibits significant difference between the green and red cluster. Taking the results of the previous section into consideration, we consider the reduction of 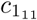 (unrecruited deacetylation) and the increase of 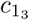 (unrecruited demethylation). The result of this reprogramming strategy is shown in Fig. 9(a), where we show that a blue cluster DERS is first transformed into a green cluster one (green square in Fig. 9(a)), and then, finally, into a red cluster DERS (red square in Fig. 9(a)). The initial blue cluster DERS has been chosen as the set with the largest value of 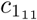, which has been shown to be a significant difference when comparing the blue cluster to the red one, and the blue cluster to the green one, leading to the idea that this property is linked to the blue cluster (idea which is reinforced because 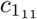 is not significant when comparing the red and the green cluster).

The efficiency of such a strategy to unlock resilient pluripotency and to encourage differentiation is shown in Fig. 9(b) where we present statistics of the differentiation time, *τ_D_*, for the original blue cluster DERS and for the corresponding reprogrammed one (two step reprogramming, *red cluster-like*). These simulations have been done for the full ER-GRN, Eqs. (17)-(18), using the hybrid multiscale simulation algorithm described in Section *Multi-scale analysis and model reduction* (and fully developed in Section S.3 of the S1 File). The resulting differentiation times for the ER-GRN with reprogrammed ER landscape are orders of magnitude smaller than those with original ER-GRN within the blue cluster DERS.

An alternative strategy, that involves changing the value of one parameter only, consists on increasing the value of 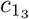 (unrecruited demethylation). Such a strategy is not obvious, since 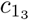 is not one of the parameters whose empirical CDF has significant differences when DERS in the red cluster are directly compared with those in the blue cluster. However, since the CDF of 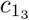 is significantly different when both the blue cluster and the red cluster are compared to the green cluster, it is conceivable that increasing 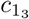 without further intervention could reprogram blue cluster DERSs. The result of this reprogramming strategy is shown in Fig. 9(a) (red diamond). Simulation results shown in Fig. 9(b) (two step reprogramming) confirm the viability of this approach. In fact, based on the statistics of the differentiation time, both strategies are virtually indistinguishable.

#### Loss of HDAC activity hinders differentiation in our ER-GRN model

Besides variability associated with cofactor heterogeneity, our model allows us to address the issue of variability regarding HME activity. HME activity is affected by both normal physiological processes, such as ageing, and pathologies such as cancer. For example, impaired activity of HDM and HDAC has been observed in relation to cancer and ageing. Here, we analyse the impact of HDM and HDAC loss of activity on the dynamics of differentiation. In particular, we simulate differentiation in our ER-GRN model to obtain statistics of the differentiation time to assess the effect of loss of HME activity. The simulations shown in this section have all been carried out using the hybrid multiscale simulation algorithm described in the S1 Appendix.

In order to clarify the effect of loss of HME activity on the ER model, we first consider the phase diagram of its mean-field limit in different situations (see [18] for details). The results are shown in Fig. S.9 of the S1 File. Figs S.9(a) and (c) show the phase diagram for two DERSs, one belonging to the red cluster (DERS1) and another, to the blue cluster (DERS2). Regarding the features of the phase diagram, the main distinction between blue cluster DERSs and red cluster DERSs, as illustrated in the examples shown in Figs. S.9(a) and (c), is that the surface occupied by the bistable region (shaded blue region) is much larger in blue cluster DERSs, because of the displacement of its lower boundary. By comparison, the bistability region of the PERSs is narrower than that of the DERSs (see Figs. S.9(b) and (d)). In particular the boundary that separates the bistable phase from the closed phase (area at the left of the blue shaded region) is displaced towards smaller HDM activity in the DERS phase diagrams.

This property suggests that a possible strategy to promote differentiation would be to decrease HDM activity, as this would drive the PERS into its closed phase whilst allowing the DERS to remain within its bistability region. In order to assess this and other scenarios, we consider a base-line scenario where the number of HMEs is exactly equal to average, i.e. *e_HDM_* = *e_HDAC_* = *Z*. We then compare different scenarios regarding the abundance of HDM and HDAC to the base-line scenario.

Contrary to what could be expected, simulation results show that the strategy of reducing HDM activity alone beyond the PERS closing boundary further hinders differentiation. As can be seen in Fig. 10(c), a decrease in HDM activity actually leads to longer differentiation time (see also [13]). Similarly, Figs. 10(a) and (b), which show statistics of the differentiation time, reveal that a decrease in both HDM and HDAC activity also leads to an increment in differentiation times, that is, this strategy fails to decrease the differentiation time below the base-line scenario. In both cases, such hindrance of differentiation is the product of the increase in the opening times (*τ*_1+_) of the DERS. This effect occurs because, as HDM and HDAC activity is reduced, the DERS is driven towards its closed-bistability boundary. Close to such a region, the DERS closed state becomes more stable and thus the corresponding *τ*_1+_ increases. By contrast, further reduction of HDAC activity moves the DERSs system closer to their open-bistable boundary, resulting in a reduction of the differentiation time. However, since the differentation times remain above those corresponding to the base line HDM and HDAC activity scenario, we conclude that loss of both HDM and HDAC activity contributes towards hindering differentiation.

**Fig 10.**
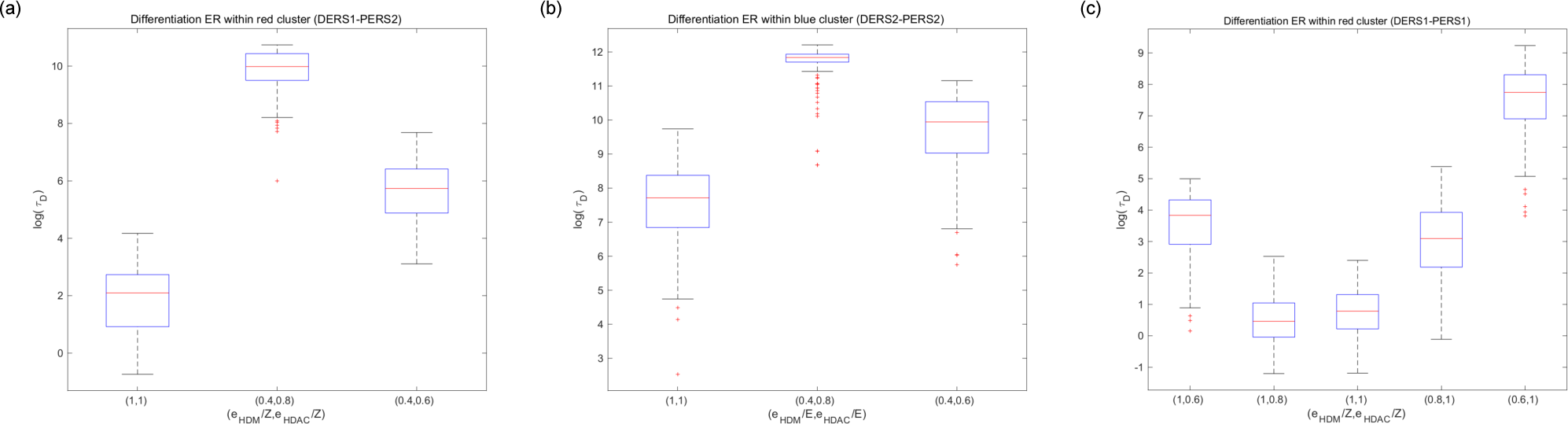
Plots showing the effect of the variation of HDM and HDAC on the statistics of the differentiation time (*τ_D_*). We consider a base-line scenario where the number of HMEs is exactly equal to average, i.e. *e_HDM_* = *e_HDAC_* = *Z*. We then compare the simulation results obtained for different scenarios regarding the abundance of HDM and HDAC to the base-line scenario, i.e. by changing the values of (*e_HDM_*,*e_HDAC_*). Parameter values: *Z* = 5 and *Y* = 15. Other parameter values given in Tables S.7, S.8, S.9 and S.10, Section S.7 of the S1 File.

## Conclusion

In this paper we have presented a model of epigenetic plasticity which has helped us to uncover some of the details and mechanisms underlying epigenetic regulation of phenotypic robustness, in particular regarding the robustness of pluripotent states. We have further uncovered how epigenetic heterogeneity regulates the decision mechanisms and kinetics driving phenotypic robustness in a stem-lock model of pathological pluripotency. Our deconstruction of epigenetic plasticity and phenotypic malleability provides crucial insights into how pathological states of permanently acquired pluripotency can be therapeutically unlocked by exploiting epigenetic heterogeneity.

We have added an ER layer to previous approaches in which cell phenotypes were associated with the attractors of complex gene regulatory systems and their robustness, with the resilience of such attractors tuned by the presence of intrinsic noise, environmental fluctuations, and other disturbances [35-43]. Our approach is based on two main pillars: namely, a framework for the generation of the ensemble of ER systems, and a multiscale asymptotic analysis-based method for model reduction of the stochastic ER-GRN model (see Section *Multi-scale analysis and model reduction*). The ensemble generation method allows the definition of epi-phenotypes based on epigenetic-regulatory modes compatible with a given state of the whole ER-GRN system [18]. We initially chose an epi-phenotype in which the ER of differentiation gene(s) (DERS) is open/active whereas the ER of pluripotency gene(s) (PERS) is closed/silent. We then used Approximate Bayesian Computation (ABC) to generate an ensemble of DERS and PERS compatible with the above-mentioned phenotype (see Section *ER-systems ensemble generation and analysis*, Fig. 3 and [18]). With such an ensemble generated, we then proceeded to evaluate its hidden, intrinsic heterogeneity in terms of the physical properties of the ER-GRN systems. This approach is closely related to the notion of neutral networks formulated to analyse systems with genotype-phenotype maps [63-65]. By making a number of assumptions regarding separation of characteristic scales, we re-scaled both the variables and the parameters of the ER-GRN system, which allowed us to discriminate the underlying separation of time scales and consequently construct an asymptotic expansion, leading to a stochastic QSSA of the system. This approximation reduces a rather complex stochastic system (Eqs. (1)-(3)) to a hybrid, 754 piece-wise deterministic Markov system (see Section *Multi-scale analysis and model reduction*). Furthermore, our model reduction procedure gives rise to an efficient and scalable, hybrid numerical method to simulate the ER-GRN system (see S1 Appendix). Although the model reduction was formulated for a GRN with an arbitrary number of mutually inhibiting genes, such a procedure is applicable to broader situations.

When analysing the behaviour of the mean-field limit of the GRN, it became apparent that, even in the simplest case considered involving a gene regulatory circuit of only two genes, the system exhibited a complex space that included several multi-stable phases. We observed a regime of tri-stability where the expected stem-lock (pluripotent) and differentiated steady-states coexist with a third state, the indecision state, in which the expression level of both genes is very low. From a developmental perspective, the latter state could serve the purpose of priming cells for differentiation, as the expression level of the pluripotency gene has decreased in a manner that could release repression upon the differentiation gene. Of note, the transitions between the different phases can be triggered by changes directly related to epigenetic regulation (i.e., cofactors of chromatin-modifying enzymes), which thereby act as bona fide molecular bridges connecting epigenetic and phenotypic plasticity by translating changes in ER states into variations of GRN states.

Having analysed the phase diagram of the GRN system and established its connection with ER (see Section *The GRN model exhibits a complex phase space, including an undecided regulatory state*), we have assessed the role of epigenetic heterogeneity in generating stem-lock pluripotent states. Such states can be viewed as examples of the so-called overly restricted epigenetic states, which present accentuated epigenetic barriers that block cell state transitions and are biologically unable to disengage self-renewal pathways [23]. The opposite situation of so-called overly permissive epigenetic states is accompanied by lowered epigenetic barriers that allow the promiscuous sampling of alternative cell states [23]. Yamanaka originally appreciated 78i the link between epigenetic heterogeneity and plasticity when aiming to explain the extremely low efficiency of somatic cell reprogramming at the population level [66]. We now know that an epigenetic predisposition to reprogramming fates exists in somatic cells and, therefore, the potential to acquire stem cell-like traits might in part reflect a pre-existing heterogeneity in cell states [26]. Furthermore, by perturbing the epigenetic state of somatic populations via inhibition of some epigenetic enzymes (e.g., the histone methyltransferase Ezh2, which catalyses repressive H3K27 methylation [67]), such heterogeneity can be harnessed to fine-tune the cellular response to reprogramming-to-pluripotency factors. Indeed, our findings support a scenario in which a sub-ensemble of ER systems with higher reprogramming potential pre-exists within the ensemble of ER systems compatible with a terminally differentiated cell state, 792 and that such a sub-ensemble could be harnessed by targeting chromatin-modifying enzymes such as HDMs and HDACs.

A careful evaluation of our ensemble of ER systems (i.e., combinations of DERSs and PERSs) concerning stem-locked systems associated with repressive ER states (see Sections *Co-factor heterogeneity gives rise to both pluripotency-locked and differentiation-primed states, Analysis of ensemble heterogeneity*, and *Ensemble-based strategies for unlocking resilient pluripotency*) has concluded that DERS heterogeneity had a stronger influence on such pluripotency-locked systems when compared with that of PERSs. Accordingly, we found that the ensemble of DERSs can be divided into three different clusters, with each one exhibiting distinct properties regarding stem locking. The so-called red cluster appeared to generate differentiation-permissive ER systems irrespective of their PERSs counterparts. By contrast, the so-called blue and green clusters contained DERSs yielding pluripotency-locked ER systems irrespective of their PERSs companions. In light of these findings, we conducted a detailed comparative analysis to uncover the underlying, statistically significant differences between DERSs within the differentiation-permissive sub-ensemble and those associated with the differentiation-repressive epigenetic states (see Section *Co-factor heterogeneity gives rise to both pluripotency-locked and differentiation-primed states*). This approach allowed us to detect which kinetic ER parameters were key to determine whether a DERS within a given system produces either permissive or repressive ER differentiation systems (see Section *Analysis of ensemble heterogeneity*). Remarkably, the elucidation of the identity of such critical regulators (see Fig. 11) would allow the formulation of strategies aimed to unlock differentiation-repressive epigenetic states by solely changing the values of such parameters (i.e., epigenetic cofactors). The feasibility of such strategies was verified by direct simulation of the ER-GRN system using our hybrid simulation method.

**Fig 11.**
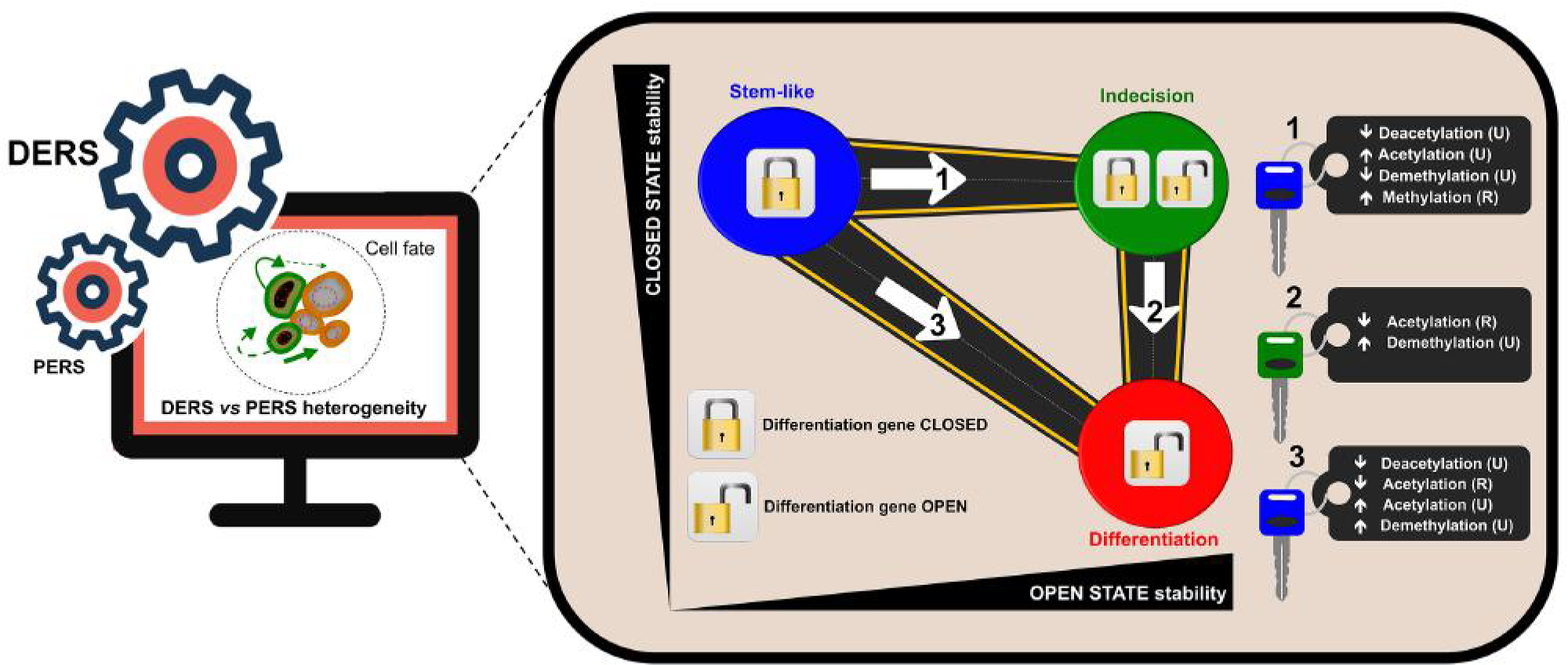
Strategies to unlock pluripotent stem-like states in ageing and cancer. Epigenetic regulation heterogeneity of differentiation genes (DERS), but not that of pluripotency genes (PERS), was predominantly in charge of the entry and exit decisions of the pluripotent stem-like states (blue). The application of the hybrid numerical method validated the likelihood of epigenetic heterogeneity-based strategies capable of unlocking and directing the transit from differentiation-refractory to differentiation-primed (red) epistates via kinetics changes in epigenetic factors. (Note: The epigenetic parameters regulating the entry into robust epi-states throughout the entire ER-GRN system revealed a regime of tri-stability in which pluripotent stem-like (blue) and differentiated (red) steady-states coexisted with a third indecisive (green) state). (R: Recruited; U: Unrecruited).

Our mathematical deconstruction of epigenetic plasticity suggests, for the first time, that epigenetic heterogeneity may underlie the predisposition of cell populations to pathological reprogramming processes that cause a permanent, locked stem-like state disabled for reparative differentiation and prone to malignant transformation. Just as the potential of single somatic cells to generate pluripotent lineages reflects a pre-existing epigenetic heterogeneity permissive for the enhancement of reprogramming fates [13,18,26], we show that ER heterogeneity could generate a subpopulation in which robustness of the pluripotent phenotype is inherently boosted. Moreover, the uncovering of the epigenetic mechanisms underpinning such stem-locked states might help in the formulation of strategies capable, for instance, of unlocking the chronic epigenetic plasticity of senescence-damaged tissues while stimulating differentiation of such stem cell-like states to successfully achieve tissue rejuvenation. As we enter a new era of therapeutic approaches to target ageing per se (e.g., senolytic agents), our current mathematical modelling and computation simulation might pave the way to incorporate new systemic strategies based on the local availability of epigenetic cofactors capable of fine-tuning the senescence-inflammatory regulation of reparative reprogramming in ageing and cancer.

## Supporting information

### S1 File. Supplemental material

#### S1 Appendix. Numerical method

In Section *Multi-scale analysis and model reduction*, we have exploited separation of time scales to formulate a QSSA whereby both the number of bound sites within the promoter of the genes and the number of enzymes and complexes molecules associated with the ER enzyme kinetics are sampled from their QSSA PDFs. Furthermore, since *S* ≫ 1, we have taken a large-*S* limit which allows us to write the dynamics of *X_i_*, i.e. the number of protein transcripts of gene *i*, in terms of an ODE perturbed by two random forcings: one associated with the random (fast) binding/unbinding dynamics and another one produced by the random ER dynamics. However, as long as we assume that *ε*_2_ = *O*(1) and *Y* = *E* ≪ *S*, further simplification is not possible and the evolution of the slow ER variables (i.e. number of positive and negative marks, and unmarked sites) need to be solved by numerical simulation of their stochastic dynamics. In spite of this, Eqs. (11)-(14) and (17)-(18) provide the reduced version of the original stochastic model which allows for a far more efficient numerical implementation of a complex ER-GRN stochastic system.

The asymptotic reduction of the full stochastic model provides the basis for a hybrid numerical method with enhanced performance with respect to the stochastic simulation algorithm (as illustrated in Figure S.4 in the S1 File). The current hybrid method is based on that formulated in [68]. The numerical method proceeds through iteration of a basic algorithm composed of the following steps:

1. Set initial conditions for the slow variables of the GRN and ER components of the system described by Eqs. (11)-(14).
2. Sample the fast variables from their QSSA PDFs conditioned to the current value of the corresponding slow variables. Their sampled values are fed in the evolution equations of the latter.
3. Consider the stochastic dynamics of the slow ER variables, Eqs. (7). These stochastic equations must be solved by numerical simulation using Gillespie’s SSA. 862 We first set the corresponding time step, δ*τ*, using the SSA.
4. Solve the ODEs for the slow variables of the GRN dynamics in the time interval [*τ*, *τ* + δ*τ*).
5. Complete the Gillespie step for the slow ER variables by choosing which elementary reaction alters the ER regulatory state and update the slow ER variables accordingly.
6. Repeat Steps through to until some stopping condition is satisfied.

If we are running the fixed time step version of the algorithm, Step needs to be done only once during the initialisation of the algorithm.

**S1 Table. The variables and parameters involved in our model formulation.**

**S2 Table. The transition rates associated with the stochastic dynamics of the GRN submodel.**

**S3 Table. The transition rates associated with the stochastic dynamics of the ER submodel.**

**S4 Table. Re-scaled GRN and ER parameters.**

## Acknowledgments

The authors would like to thank Heather Harrington and Mariano Beguerisse for helpful discussions and advise. This work is supported by a grant of the Obra Social La Caixa Foundation on *Collaborative Mathematics* awarded to the Centre de Recerca Matematica. The authors have been partially funded by the CERCA Programme of the Generalitat de Catalunya. E.C. is the recipient of a Sara Borrell post-doctoral contract (CD15/00033, Ministerio de Sanidad y Consumo, Fondo de Investigation Sanitaria, Spain). N.F-B. and T.A. acknowledge MINECO and AGAUR for funding under grants MTM2015-71509-C2-1-R and 2014SGR1307. T.A. acknowledges support from MINECO for funding awarded to the Barcelona Graduate School of Mathematics under the “Maria de Maeztu” programme, grant number MDM-2014-0445. R.P-C also acknowledges the UCL Mathematics Clifford Fellowship. This work was supported by grants from MINECO (SAF2016-80639-P) and AGAUR (2014 SGR229) to Javier A. Menendez.

